# Allelic turnover and dominance reversal at a single-gene balanced polymorphism controlling heterodichogamous flowering in wingnuts (Juglandaceae)

**DOI:** 10.1101/2025.03.10.642504

**Authors:** Jeffrey S. Groh, Gracie Ackerman, Michael Wenzel, Graham Coop

**Affiliations:** Department of Evolution and Ecology, University of California, Davis; Center for Population Biology, University of California, Davis; Sonoma Botanical Garden, Glen Ellen, California

## Abstract

Angiosperms have evolved a wide variety of spatial and temporal developmental mechanisms to limit sexual interference and inbreeding. In heterodichogamy, two hermaphroditic morphs exploit distinct temporal reproductive niches by alternating phases of male and female flowering, promoting disassortative mating. This system is widespread within Juglandaceae, and known to be controlled by ancient balanced polymorphisms in walnuts (*Juglans*) and hickories (*Carya*). Here we identify distinct inheritance mechanisms controlling heterodichogamy in two separate Juglandaceae genera, the wingnuts (*Pterocarya* and *Cyclocarya*). We first document the occurrence of heterodichogamy in *Pterocarya* and map its genomic basis to haplotypes overlapping a single candidate gene in the *FANTASTIC FOUR* superfamily (*GFAFL*). These haplotypes segregate throughout the entire genus and the dominant haplotype controls female-first flowering. We show heterodichogamy in the sister genus *Cyclocarya* is associated with a distinct pair of ancient haplotypes at the same locus in both diploids and tetraploids, but with a dominant allele controlling male-first flowering. We infer a well-resolved fossil-calibrated phylogeny of Juglandaceae and date the divergence of the *Pterocarya* and *Cyclocarya* haplotypes to 51 and 44 million years ago, more recent than the divergence of these genera. In *Pterocarya* female-first heterozygotes, the dominant haplotype is associated with allele-specific suppression of the recessive copy of *GFAFL* during early male flower development. In *Cyclocarya* male-first heterozygotes, the dominant haplotype is associated with allele-specific activation of the recessive copy of *GFAFL* during male flower development, while the dominant copy itself shows higher expression in female flowers. We propose a model for the evolution of reciprocal sex matching in heterodichogamy through the combination of *cis*-regulatory divergence and allelic interactions involving ‘fast’ and ‘slow’ alleles at a single gene regulating flowering time. The non-independent expression of alleles in both systems is reminiscent of *trans*-sensing phenomena in other systems and suggests a mechanism mediated by DNA homology or an RNA intermediary. Notably, the dominant haplotypes in both genera show parallel architecture with a hemizygous region containing tandem duplicates of the same 1 kb motif downstream of the transcribed region of *GFAFL* which may be linked with such a mechanism. Our findings shed light on the molecular basis of heterodichogamy and contribute to an emerging view that diverse genetic pathways can be co-opted during its evolution and facilitate turnover in its genetic control.

## Introduction

The timing of reproduction is a critical aspect of fitness and is shaped by adaptive evolution (Hall and Willis 2006; Franks *et al.* 2007; Gaudinier and Blackman 2020). In hermaphroditic organisms which can produce both types of dimorphic gametes or gametophytes, the optimal timing of their development, i.e. reproduction through ‘male’ vs. ‘female’ pathways, may differ as a result of resource trade-offs, sex-specific changes in fecundity through time, and costs associated with self-fertilization and gamete interference. These factors are thought to underlie the evolution of diversity in sex-specific temporal mating strategies such as sequential hermaphroditism in animals and dichogamy (temporal separation of male and female function) in flowering plants (Warner 1975; Munday *et al.* 2006; Lloyd and Webb 1986; Routley *et al.* 2004).

Simultaneously with individual-level trade-offs, sex-specific temporal strategies are also shaped by sex-ratio selection acting within a population (Leigh 1976; Brunet and Charlesworth 1995). Heterodichogamy, found in at least 13 angiosperm families, refers to a stable dimorphism for the temporal order of staminate vs pistillate flowering (i.e. male and female) (Renner 2001; Endress 2020). Protandrous (male-first) individuals release pollen prior to their stigmas being receptive, whereas protogynous (female-first) individuals show the complementary pattern. Pollen dispersal of one morph thus occurs synchronously with stigma receptivity of the other. This promotes disassortative outcrossing (Bai *et al.* 2007), such that the rare mating type is favored and a 1:1 ratio of two morphs is expected at equilibrium (Gleeson 1982). This mating system is thus analogous to dioecy and heterostyly, in which discrete floral morphs are maintained by the negative frequency dependence of disassortative mating between morphs (Fisher 1930; Ganders 1979). Heterodichogamy is suggested to facilitate the evolutionary transition from hermaphroditism to dioecy (Pendleton *et al.* 2000; Pannell and Verdú 2006), though itself can remain stable over vast evolutionary time scales (Groh *et al.* 2025). As with dioecy and heterostyly, heterodichogamy appears to typically be controlled by single-locus mechanisms involving hemizygous functional regions and sometimes recombination suppression (Groh *et al.* 2025; Zhao *et al.* 2025), revealing generalities on the genomic causes and consequences of different mating systems. However, the mechanisms of gene regulation controlling heterodichogamy remain poorly understood, and may show unique features.

A classic example of heterodichogamy is its widespread occurrence in the walnut family (Juglandaceae) (Delpino 1874; Darwin 1877; Fukuhara and Tokumaru 2014; Mao *et al.* 2016). This system is present in the many species of walnuts (*Juglans*) and hickories (*Carya*, e.g. pecan) examined to date, and in both genera it is controlled by Mendelian segregation of a dominant allele for protogyny (G dominant, g recessive) (Gleeson 1982; Thompson and Romberg 1985). In our previous work, we investigated the molecular basis and evolutionary history of heterodichogamy within Juglandaceae, and found that two distinct genomic regions involving different genes control the mating system in walnuts (*Juglans*) and hickories (*Carya*, e.g. pecan) (Groh *et al.* 2025). The pecan system resembles a supergene, with a region of reduced recombination spanning numerous flowering time candidate genes, whereas the walnut system involves *cis*-regulatory divergence of a single candidate gene for flowering time. Both systems are ancient (ca. 40-50 Mya), yet the loci span relatively narrow genomic regions (⇠20-400 kb) unlike heteromorphic sex chromosomes. Within Juglandaceae, heterodichogamy has been reported in two additional genera (*Cyclocarya* and Platy*Carya*); however, we previously showed the genetic basis of the mating system in these genera was not shared with either walnuts or hickories (Groh *et al.* 2025), suggesting that additional genetic mechanisms control heterodichogamy in these other genera.

Here we focus on the flowering behavior of two Juglandaceae genera known as wingnuts - (*Pterocarya*), in which heterodichogamy had not been previously documented to our knowledge, and the sister genus *Cyclocarya* (wheel wingnut). Wingnuts are so named for their fruits (Fig. 1d), which are suited for wind dispersal, unlike the animal-dispersed husked seeds of *Juglans* and *Carya*. As in *Juglans* and *Carya*, unisexual flowers of *Pterocarya* and *Cyclocarya* are borne in separate male and female catkin inflorescences (Fig. 1a,b), or rarely in the same inflorescence (Fig. S1, Pigg *et al.* 2023) and pollen is wind-dispersed. Macrofossils of *Cyclocarya* appear in the early Paleocene (Lyson *et al.* 2019), whereas fossil *Pterocarya* fruits are not known prior to the Oligocene (Yan *et al.* 2024). The fossil diversity and distribution of these genera throughout Neogene sediments of North America, Europe and Asia indicate that wingnuts were once considerably more diverse and geographically widespread than today (Manchester 1987). *Pterocarya* remains present in the fossil record of North America and Europe through the Pliocene, and appears to have gone extinct in these regions during the Pleistocene glaciations (Song *et al.* 2021; Stults *et al.* 2022). The current distribution is accordingly thought to reflect locations of glacial refugia; of the six recognized species, five are restricted to East Asia and a single species (Caucasian wingnut, P. fraxinifolia) is endemic to the Caucasus region. *P. stenoptera* is the most wide-spread species, being found across China, and it is used in urban landscaping in both Europe and North America. *Cyclocarya* contains a single modern representative native to China (C. paliurus), which consists of both diploid and autotetraploid lineages that diverged ⇠570,000 years ago (Qu *et al.* 2023b; Yu *et al.* 2023).

**Figure 1:**
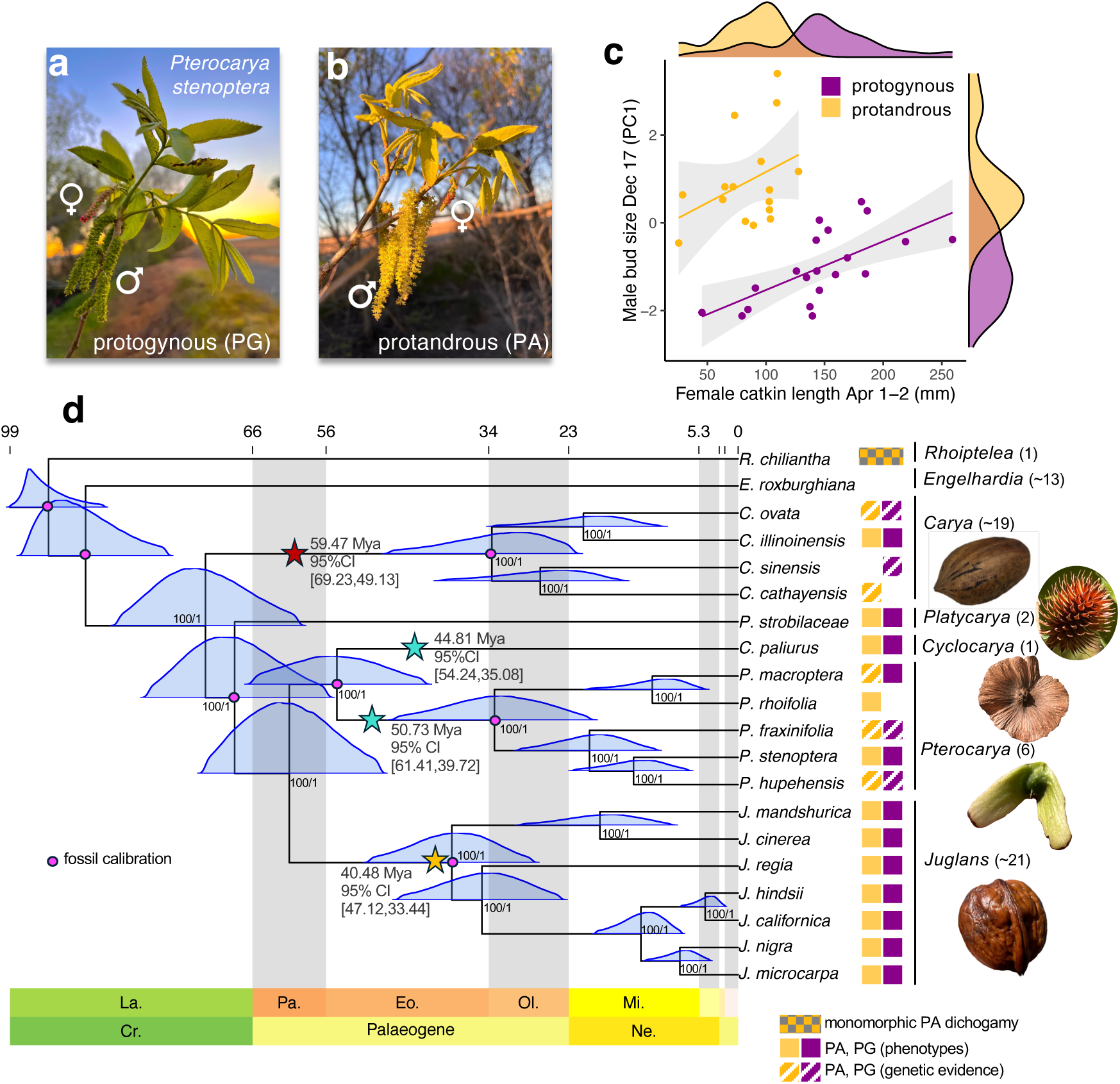
Heterodichogamy in **Pterocarya** and the phylogeny of Juglandaceae. **a,b)** Individuals of **Pterocarya* stenoptera* show either protogynous (a) or protandrous (b) dichogamy. In (a), mature stigmas (pink) are visible on the pistillate inflorescence, while anthers on the staminate inflorescence have not yet released pollen. In (b), anthers have released pollen, and pistillate flowers are beginning to reach maturity. **c)** Protandrous (gold) and protogynous (purple) individuals show reversed dimorphism for developmental stage of staminate and pistillate inflorescences. Density plots for each inflorescence type are shown at top and right. Developmental size of staminate and pistillate inflorescences are overall negatively correlated due to the dimorphism, but are positively correlated within each types. **d)** Fossil-calibrated phylogeny of Juglandaceae. Time (in units of MYA) shown at top with abbreviations of geological periods and epochs at bottom. Full posterior distributions on node ages are shown in blue. Node labels indicate bootstrap support from a maximum likelihood analysis and local posterior probability values from ASTRAL. Pink circles indicate nodes with priors specified on the basis of fossil information (see Supplemtary Text 1 and Fig. S3). Colored rectangles at right indicate the type of dichogamy observed in each taxon. Absence of colored rectangles means dichogamy types have not been reported. Full genus names are written at right with species numbers in parentheses and a representative seed structure. Colored stars indicate the origins of separate genetic systems controlling heterodichogamy in Juglandaceae, with estimates and credible intervals.

We combined denovo chromosome-level assemblies, resequencing of phenotyped flowering individuals, and gene expression data in order to characterize the evolutionary history and molecular basis of heterodichogamy in wingnuts. Our study identifies a novel molecular mechanism underpinning this mating system in *Pterocarya* and parallel use of the same gene in *Cyclocarya*, but with a different set of alleles and a reversed dominance pattern. Our results inform a model for the evolution of heterodichogamy and further contribute to a picture of dynamic genetic control of heterodichogamy in Juglandaceae, despite the evolutionary stability of the mating system and conserved inheritance mechanisms within genera.

## Results

### **Pterocarya** exhibits heterodichogamous flowering

We studied flowering of dozens of *Pterocarya* stenoptera individuals from germplasm collections and urban plantings and ‘volunteer’ trees to assess the relative timing of development of male and female flowering structures. Individuals showed clear dichogamy of two types. Protandrous (male-first) individuals (Fig. 1a) released pollen prior to stigma maturity, while protogynous (female-first) individuals showed mature stigmas prior to pollen release (Fig. 1b). Among plants at the same locality, these timing differences resulted in the reciprocal matching of pollen release of one morph with stigma receptivity of the other.

The offset timing of staminate and pistillate flowering of the two morphs was reflected in quantitative measurements of inflorescence size (Fig. 1c). Female inflorescences, which progressively lengthen throughout flowering and post-fertilization, are longer in protogynous individuals mid-season. On the other hand, male inflorescences, which begin development in the season prior to flowering, enter dormancy at a larger size in protandrous individuals (Fig. 1c), a pattern which has also been observed in heterodichogamous *Cyclocarya* (Chen *et al.* 2019), *Juglans* (Luza and Polito 1988; Groh *et al.* 2025) and *Carya* (Rhein *et al.* 2023).

Protandrous and protogynous morphs were common across sites and segregate among maternal siblings from mother trees with different collection sources (Fig. S2). Among all flowering individuals examined across sampling localities, we identified 36 protandrous individuals, 36 and protogynous individuals, and 17 individuals whose dichogamy type could not be assessed due to limited observation.

We observed flowering in a single individual from each of two sister species, P. macroptera and P. rhoifolia. The P. macroptera individual showed protogynous flowering, while the P. rhoifolia individual exhibited protandrous flowering. Segregating dichogamy types in two clades spanning the deepest divergence within *Pterocarya* (Fig. 1d), together with the prevalence of the mating system throughout Juglandaceae, strongly suggest that heterodichogamy occurs in all species of *Pterocarya*.

### Genome assembly and annotation

To facilitate investigation into the molecular basis and evolutionary history of heterodichogamy in wingnuts, we generated haplotype-resolved chromosome-scale genome assemblies from three species of *Pterocarya* and from their sister group *Cyclocarya* paliurus using PacBio HiFi sequencing and scaffolding to Hi-C aided long-read assemblies from the same genera (Zhang *et al.* 2024d; Qu *et al.* 2023a). *Pterocarya* assembly samples originated from a single flowering individual each of *P. stenoptera*, *P. macroptera* var. insignis (both protogynous flowering types) and a non-reproductive P. fraxinifolia individual of unknown flowering type. The diploid C. paliurus individual from Sonoma Botanical Garden (formerly Quarryhill Botanical Garden) used for assembly showed protogynous flowering and is one of two known trees of this species established in North America (Sutton 2019).

Our assemblies are highly contiguous; for example, haplotype 1 of *P. stenoptera* has a contig N50 of 32.14 Mb, with 97.9% of the 551.2 Mb assembled genome contained within 16 chromosome-scale scaffolds (the expected number from cytological counts, Oginuma 1999) and an average of 2.38 contigs per chromosome. After masking repetitive sequences (54% of the genome), we annotated our *P. stenoptera* assemblies leveraging extrinsic evidence from protein homology and RNA-seq datasets from 7 tissues (data from this study and Zhang *et al.* 2024d), resulting in 30,681 gene models for haplotype 1 and a BUSCO completeness score of 98.4%. Other assemblies showed similarly favorable metrics (Table S1).

### A time-calibrated phylogeny of Juglandaceae

To establish the age and distribution of Juglandaceae heterodichogamy systems in a common phylogenetic framework, we constructed a robust fossil-calibrated phylogeny by combining our de-novo assemblies with whole genome sequence data spanning the diversity of Juglandaceae (representing all recognized tribes and 7 of the 9 extant genera sensu APG IV 2016). Using a dataset of 5,089 single copy orthologs containing 443,387 parsimony-informative sites, we recovered a well-resolved species tree, with both gene-tree summary approaches (ASTRAL, Zhang *et al.* 2018) and concatenation + maximum likelihood showing strong support for the same topology (Fig. 1d). This topology is largely in agreement with previous molecular phylogenies of Juglandaceae that included broad taxon sampling and genome-scale nuclear datasets (Mu *et al.* 2020; Zhou *et al.* 2021; Ding *et al.* 2023), and resolves *Carya* rather than Platy*Carya* as the outgroup within subfamily Juglandoideae (but see Ding *et al.* 2023).

We time-calibrated the phylogeny in a Bayesian framework (using MCMCTree, Reis and Yang 2011) by combining fossil data to specify time priors (Supplementary Text 1, Fig. S3) with likelihood under a relaxed-clock substitution model. The time-calibrated phylogeny (Fig. 1d, Table S2) supports an origin of crown Juglandaceae in the Late Cretaceous (94 Mya, 95% Credible Interval 86-99), with a subsequent divergence of the various tribes occurring near the K-Pg boundary. We infer an early Oligocene or late Eocene crown age of *Pterocarya* (31 MYBP, 95% CI 20-47) and subsequent radiation during the Miocene, with relationships supporting reciprocal monophyly of intrageneric sections *Pterocarya* (*P. stenoptera*, P. hupehensis, P. fraxinifolia) and Platyptera (*P. macroptera*, P. rhoifolia) (Manning 1978).

Finally, we note that the *Pterocarya*-*Cyclocarya* divergence is estimated at 55 Mya (95% CI 43-66), despite specifying a prior strongly favoring a minimum divergence of 65 Mya on the basis of early Paleocene *Cyclocarya* fossils (Lyson *et al.* 2019) (Fig. S4). This signal in the molecular data favoring a younger divergence is consistent with the lack of definitive fossils of *Pterocarya* prior to the Oligocene (Manchester 1987) (30 million years after the earliest *Cyclocarya* fossils), and suggests that *Pterocarya* could have evolved from an ancestor with *Cyclocarya*-like fruit morphology.

### Ancient haplotypes at a single locus control heterodichogamy throughout **Pterocarya**

To identify the genetic basis of dichogamy type within *P. stenoptera*, we resequenced 20 individuals (mean depth 27x), combined with existing data from 10 additional individuals (Groh *et al.* 2025), with confidently assigned dichogamy phenotypes. Our mapping population consists of a mixture of wild-derived seed collections and recently wild-derived material from several sources (Table S3) and contains multiple sets of maternal siblings segregating for dichogamy type (Fig. S2).

Using Genome-Wide Association (GWAS), we identified a single clear association peak on chromosome 11 (Fig. 2a) away from the centromere, consistent with a single-locus Mendelian inheritance mechanism. This region is not syntenic with *Juglans* chromosome 11 (Fig. S5) and does not contain homologs of any of the genes present in the *Juglans* or *Carya* heterodichogamy loci. The association peak is contained within single assembly contigs in each *P. stenoptera* haplotype assembly. The peak contains a set of SNPs perfectly predictive of dichogamy type spanning a 65.5 kb region in haplotype 2 of the assembly (Fig. 2b) (vs. 77.2 kb in haplotype 1, Fig. S6). We refer to this as the G-locus. Both assembled haplotypes contain reciprocal orthologs of a gene within the FANTASTIC-FOUR (FAF) superfamily (red, Fig. 2b), which we return to below.

**Figure 2:**
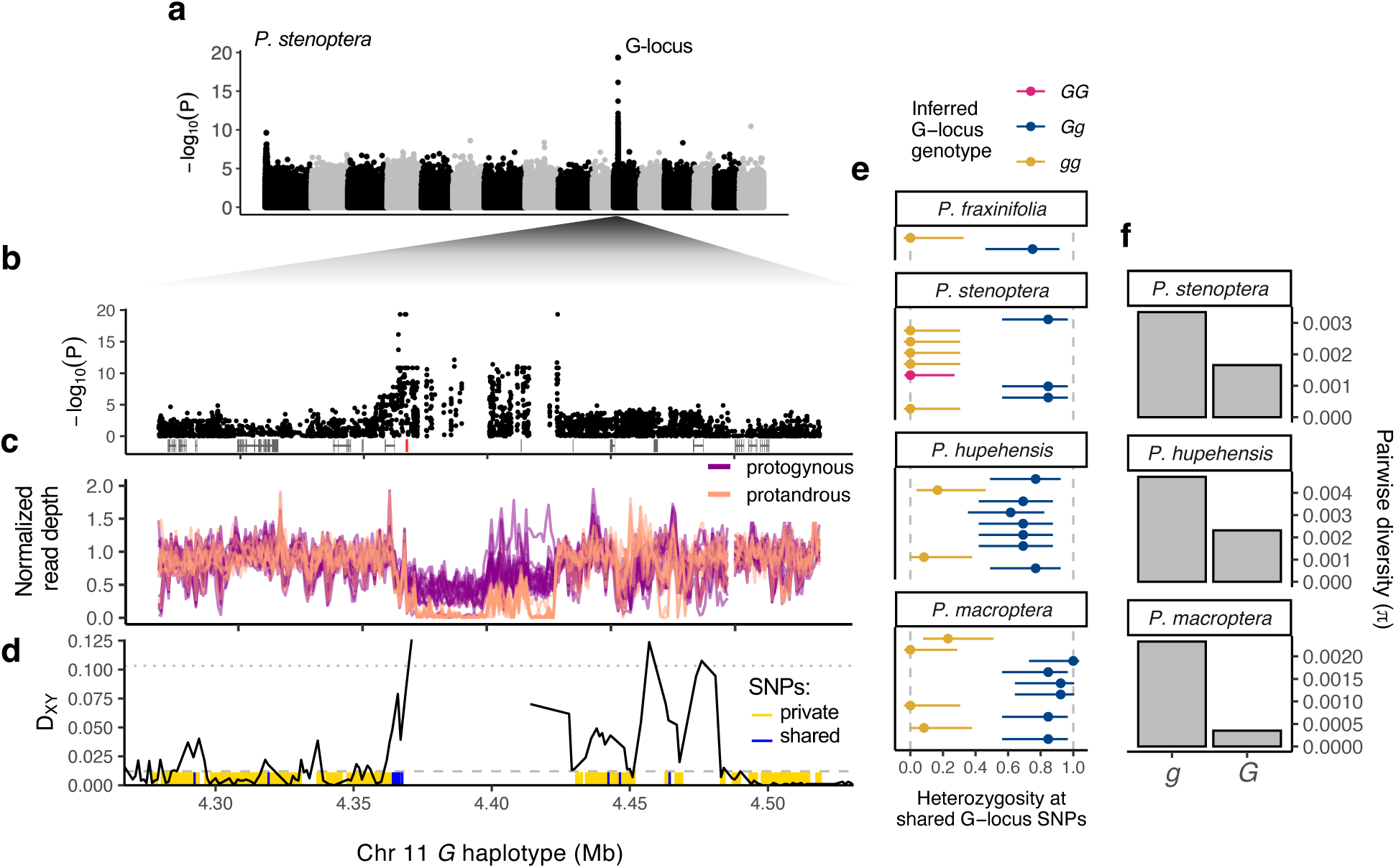
A single genetic locus for heterodichogamy in **Pterocarya**. **a)** GWAS for dichogamy type in 30 individuals of *P. stenopera* reveals a single clear peak on chromosome 11 (shown for reads aligned to haplotype 2 which contains the dominant allele). Chromosomes are arranged in increasing number, alternating black and gray. **b)** Zoomed in view of the GWAS peak (G-locus). Predicted gene models are shown at bottom with a *FAF*-like gene (*GFAFL*) in red. **c)** Normalized average read depth in 1 kb windows across the G-locus. Lines are colored by each individual’s dichogamy type. **d)** Nucleotide divergence in 1 kb windows across the region. Dashed and dotted lines show the genome-wide mean and 99% quantile respectively. Colored tick marks at bottom show positions of SNPs from a comparison of *P. stenoptera* and *P. macroptera* haplotypes where alleles group by species (gold) or are shared across species and group by haplotype (blue). **e)** Individuals in four **Pterocarya** species from published datasets (Geng *et al*. 2024; Groh *et al*. 2025) segregate for the shared SNP haplotypes identified in (d) between *P. stenoptera* and *P. macroptera*. Bars show 95% confidence intervals of the proportion. **f)** Diversity is lower in three species on the *G* haplotype compared to the *g* haplotype.

Genotypes at these SNPs are consistent with a dominant allele (G) determining protogyny, and indicate that our haplotype assemblies 1 and 2 represent the recessive (g) and dominant (G) alleles, respectively. In our GWAS sample all 15 protandrous individuals are gg homozygotes, and of the 15 protogynous individuals 14 are Gg heterozygotes while one is a GG homozygote. This deficit of homoygotes for the dominant allele is expected to result under disassortative mating. We tested this hypothesis by combining our data with published data from additional P. stenoptera individuals sampled from its current natural range (Geng *et al.* 2024). Consistent with disassortative mating between morphs, we identified only two dominant allele homozygotes (out of 40), a significant deviation away from the Hardy-Weinberg expectation under random mating (P= 1.27 ⇥ 10^-8^, *x*^2^ = 32.38, d.f.=1).

Patterns of read depth across this region showed clear stratification by phenotype (Fig. 2c). Protogynous heterozygotes appear hemizygous for a set of indels with respect to both assembled haplotypes, whereas protandrous individuals lack read depth across the entirety of the G haplotype but show typical read depth across the g haplotype (Fig. S6). These patterns point to structural differences and/or high divergence between the two haplotypes. Indeed, compared to the genome wide average level of polymorphism per base pair (⇡ = 0.008), the divergence between haplotypes for alignable sequence is exceptionally high in this region (D*_xy_* = 0.126), falling in the top 1% of windowed 1 kb values genome wide (Fig. 2d).

The high divergence across the G-locus suggests that it could be an ancient balanced polymorphism. We leveraged our haplotype assemblies from the protogynous *P. macroptera* individual and identified a high concentration of *trans*-species polymorphic SNPs at the GWAS peak (blue, Fig. 2d), whereas SNPs immediately adjacent to the GWAS peak group by species (yellow, Fig. 2d). We next examined heterozygosity at this set of *trans*-species polymorphic SNPs in published whole genome resequencing from multiple individuals each of three species (P. stenoptera, *P. macroptera*, and P. hupehensis) (Geng *et al.* 2024). Consistent with these haplotypes segregating in all three species, we find multiple individuals within each species that are highly heterozygous at this set of SNPs and others that show low heterozygosity (Fig. 2e). The phenotypes predicted by these genotypes do not significantly deviate from the null expectation of a 1:1 morph ratio in each species, consistent with disassortative mating maintaining the polymorphism within populations. We also detected shared polymorphism at these SNPs in one of two previously sequenced individuals of P. fraxinifolia (Groh *et al.* 2025) (Fig. 2e), and confirmed that a previously sequenced protandrous P. rhoifolia individual lacked shared polymorphism at these SNPs. We used estimates of species divergence times from the genome-wide phylogeny to calibrate a molecular clock for this region (see Methods), inferring that the *Pterocarya* haplotypes diverged 51 Mya (95% CI: 40-61), in the ancestor of extant *Pterocarya*.

The region tightly linked to the G allele is predicted to share evolutionary features with Y-linked sex determining regions. As disassortative mating maintains the G haplotype at lower frequency compared to the g haplotype, under a neutral model we expect a 3-fold reduction in diversity among G vs. g haplotypes with complete disassortative mating. In addition, the G haplotype may rarely recombine. The physical extent of conserved *trans*-species polymorphism indicates a lack of historical recombination between G and g haplotypes, and G may also rarely recombine with itself because GG homozygotes are rare. Thus, effects of hitchhiking and/or Muller’s ratchet will be felt stronger on G haplotypes. Using phased resequencing data across the coding sequence of the FAF-like gene, we find support for these predictions, with G haplotypes showing a 2-6.6-fold reduction in several species (Fig. 2f). Y-linked regions and other permanently heterozyogous supergenes often accumulate repetitive sequences at higher rates than autosomally-inherited regions (Bachtrog 2013; Stolle *et al.* 2019; Akagi *et al.* 2025; Moraga *et al.* 2025). Despite the fact that the G haplotype is nearly always heterozygous, the G and g haplotypes are of similar lengths in both *Pterocarya* species. G-locus haplotypes are only modestly enriched in the percentage of repeat content compared to their local genomic context, and this enrichment is not stronger for the G haplotype (Fig. S7). The lack of considerable expansion of the dominant haplotype suggests that degenerative impacts of heterodichogamous mating on molecular variation are limited in *Pterocarya*.

### Allele-specific regulation of a single flowering gene at the **Pterocarya** G-locus

The region of *trans*-species G-locus SNPs coincides closely with the transcribed region of a *FANTASTIC FOUR* (FAF) domain-containing gene, which we refer to here as G-LOCUS-FAF-LIKE (GFAFL) (Fig. 3a). The FAF domain (Fig. S8) is a plant-specific domain of unknown molecular function named for a group of four Arabidopsis genes identified as regulators of shoot meristem size (Wahl *et al.* 2010). Other FAF domain-containing (FAF-like) genes have since been found to regulate diverse developmental processes including abscisic acid signaling, abiotic stress tolerance, endosperm starch metabolism (Wang *et al.* 2018; Lim *et al.* 2022; Zhang *et al.* 2019), and flowering time (Zhang *et al.* 2024a). We identified FAF-like paralogs in several embryophytes and find that all previously functionally characterized members of this group are distant paralogs to GFAFL (Fig. 3c). Nonetheless, orthologs of GFAFL in *Cyclocarya*, *Juglans* and Arabidopsis show flower- or pollen-specific expression (Figs. S9, S10) (Chakraborty *et al.* 2016; Hardwood Genomics Team 2015; Schmid *et al.* 2005; Klepikova *et al.* 2016), supporting a conserved role of GFAFL in regulating flowering.

**Figure 3:**
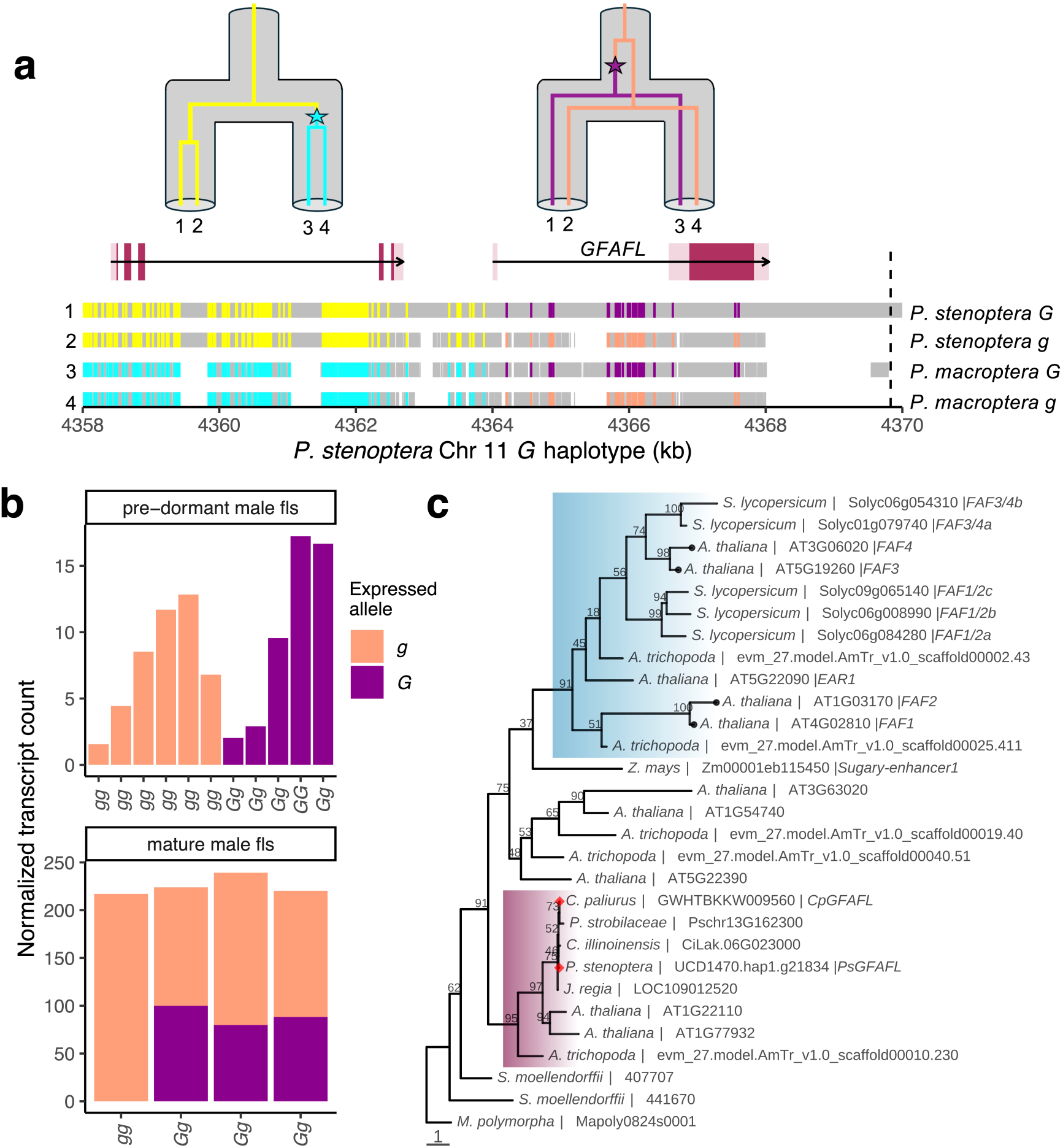
*GFAFL* is a candidate gene for the regulation of heterodichogamous flowering in **Pterocarya**. **a)** Outside the region of copy number variation, *trans*-species polymorphic SNPs at the **Pterocarya** G-locus localize to the transcribed region of *GFAFL*. Each horizontal bar represents a haplotype alignment to the *P. stenoptera G* reference, with gray portions showing aligned regions. Positions with colored tick marks are biallelic SNPs (singletons not shown) with the alleles in two different colors. Above, we show the implied tree sequence, noting that across sites either allele may be the derived mutation. The dotted vertical line at right indicates the start position of the region homologous to the repeats contained within the dominant haplotype (*G-0*). **b)** *GFAFL* shows allele-specific expression in male flowers. Each bar represents a unique individual, genotypes are shown at bottom as determined from genomic sequence data. **c)** Maximum likelihood phylogeny of *FAF*-like genes in a selection of embryophytes from a codon-based alignment of the *FAF* domain, rooted using a single copy gene in *Marchantia*. The subclade highlighted in blue contains the canonical Arabidopsis **FANTASTIC FOUR** genes (black points at tips). *GFAFL* is a member of a different subclade (maroon) which we infer is related to the previous via gene duplication in the ancestor of angiosperms. Copies in **Pterocarya** and **Cyclocarya** are indicated with red diamonds. Alignment across a larger coding region in this subclade reveals *GFAFL* shows closer homology to the Arabidopsis gene *AT1G22110* (Fig. S8). Node labels give bootstrap support. Branch lengths represent the number of substitutions per codon site.

In the coding region of GFAFL we see only one nonsynonymous difference between haplotypes conserved across P. stenoptera and *P. macroptera*, a nonconservative K to N replacement in the recessive haplotype lineage at an otherwise conserved codon site (Fig. S8). Within species, the haplotypes show greater divergence across several conserved regions; the *P. macroptera* haplotypes are ⇠2.5% diverged at the level of amino acid sequence (*^dN^*/*dS* = 0.58), while the P. stenoptera haplotypes are (⇠8%) diverged in amino acid sequence (*^dN^*/*dS* = 0.65). Using phased resequencing data, we did not detect any signal of adaptative molecular evolution as there was no excess of nonsynonymous vs. synonymous substitutions between the haplotypes using nonsynonymous and synonymous polymorphism among g haplotypes as a control (MacDonald-Kreitman test).

To investigate the role of gene expression regulation in the developmental basis of heterodichogamy, we performed RNA-seq from both male and female flower tissues of both morphs collected on the same day during flowering (March 20^th^), as well as from pre-dormant male catkin buds (on Aug. 2^nd^) in the same year. These three tissues show distinct genome-wide expression patterns across biological replicates, and we see a signal of transcriptome-wide differentiation between morphs within mature flowering tissues (Fig. S11). Examining RNA-seq read alignments from both reproductive and vegetative tissues (data from Groh *et al.* 2025; Zhang *et al.* 2024d), we found that GFAFL is the only gene in the G-locus region with (1) unequivocal evidence of expression in floral tissue and (2) conserved presence across multiple G-locus haplotypes of *Pterocarya* (see Supplementary Text 2).

In P. stenoptera, we found expression of GFAFL in male catkins and no other tissues, with a large increase in expression between the bud stage and floral maturity (Fig. 3b). Neither overall expression levels nor splicing patterns of the gene appear to differ between morphs. However, we identified patterns of allele-specific expression using G-locus SNPs within RNA-seq reads. In early development of male flowers, heterozygotes for the G-locus show a strong bias toward expression of the G allele of GFAFL (P<0.01, x^2^ = 7.11, d.f.=1). Moreover, expression of g in heterozygotes is entirely lacking, which is significantly less than what would be expected compared to half the expression level seen in gg homozygotes (P<0.001, x^2^ = 15.11, d.f.=1). The highest expression of G was seen in the only GG homozygote; which could suggest a dosage-sensitive effect, but this effect is weak and not statistically significant. In mature male flowers, the pattern of allele-specific expression in heterozygotes appears to weaken or even reverse, and the level of expression of g is not reduced compared to its expectation from the gg homozygote.

In the early stages of male flower development, despite the lack of g expression of GFAFL in heterozygotes, we see clear expression of the G copy (Fig. 3b). These facts suggest a mechanism of allele-specific *trans*-silencing of the g copy of GFAFL by the G haplotype.

### Heterodichogamy in **Cyclocarya** is also associated with *GFAFL* but with reversed dominance

The protogynous C. paliurus individual we used for assembly is not heterozygous for the two *Pterocarya* dichogamy haplotypes described above, nor is it homozygous for the dominant haplotype seen in *Pterocarya*. This implied there must be a distinct inheritance mechanism for heterodichogamy in *Cyclocarya*. Remarkably, we find evidence that a distinct pair of haplotypes at the same locus with reversed dominance control heterodichogamy in both the diploid and tetraploid lineages of *Cyclocarya*.

First, we detected flower-specific expression of a highly divergent copy of GFAFL in published expression data from C. paliurus (Zhang *et al.* 2024b), which was not found in either of the assembled haplotypes or raw sequencing reads from our protogynous C. paliurus individual, nor in high-coverage sequencing reads from a different known protogynous individual (Qu *et al.* 2023a). We identified that this GFAFL transcript maps to coding sequence at the syntenic location (hereafter, the G2 allele) in assemblies from both diploid and tetraploid protandrous individuals (Qu *et al.* 2023a). In the diploid consensus assembly, G2 appears as a tandem paralog to its counterpart (G1), however closer inspection revealed this assembly to be chimeric, and the diploid individual is heterozygous for two highly diverged orthologous alleles (Fig. S13). In the haplotype-resolved assembly of a protandrous tetraploid, GFAFL is single-copy in each of four assembled haplotypes at the syntenic location, with one G2 and three G1 alleles (Fig. 4a). In a separate RNA-seq dataset (Chen *et al.* 2019), we detected expression of G2 only in labeled protandrous samples, and not in protogynous samples (representing 9 individuals of each type).

**Figure 4:**
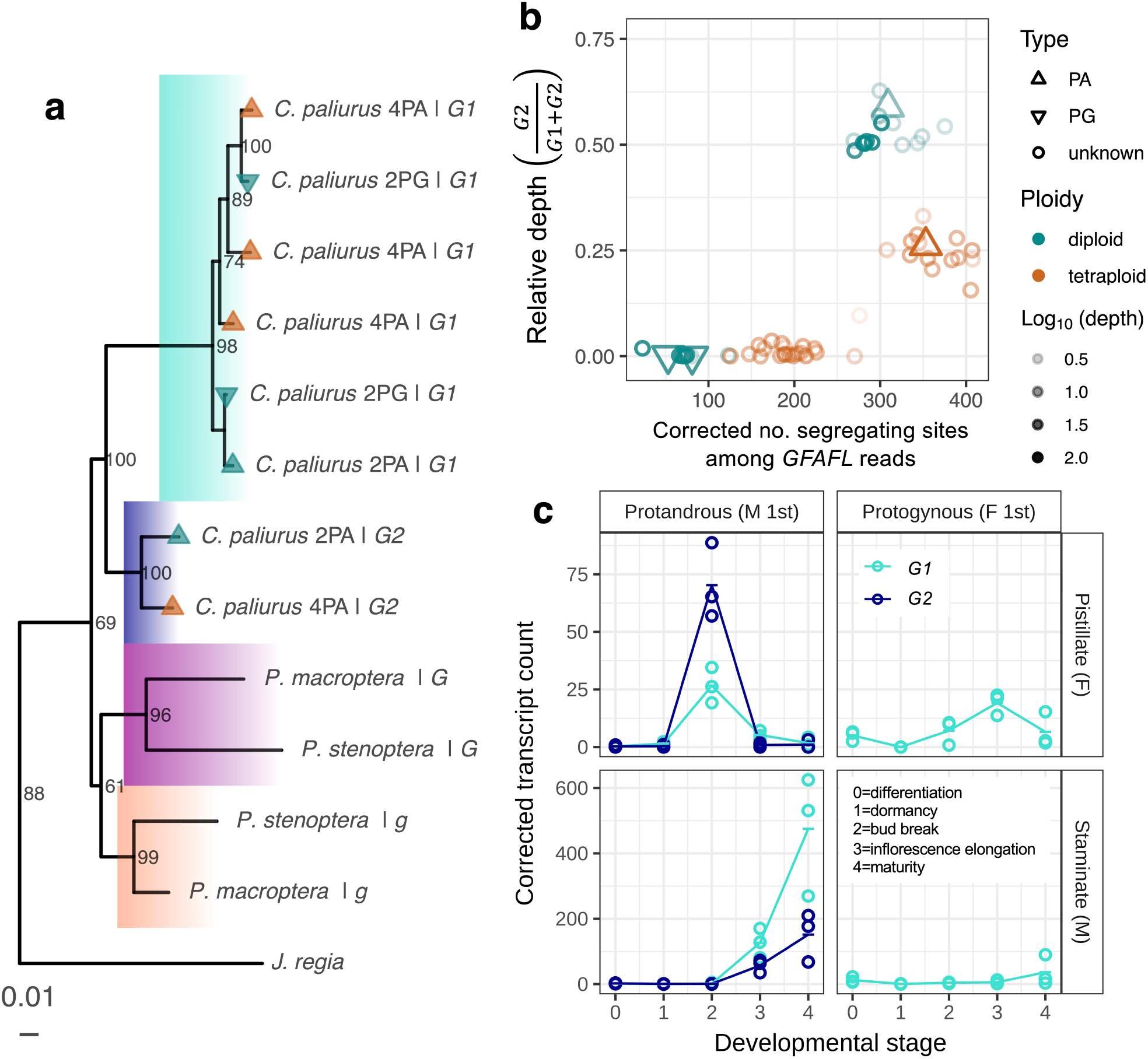
**a)** Phylogeny of the *GFAFL* intron across **Pterocarya** and **Cyclocarya** assemblies, rooted with *Carya illinoinensis* as an outgroup (not shown). Tip shapes and colors indicate the flowering type and ploidy of the individual from which each *GFAFL* copy derives. **b)** Segregation of the *G2* allele in both diploid and tetraploid lineages of *C. paliurus*. Each point represents a unique individual. The y-axis shows relative read depth across alternate alleles of the *GFAFL* transcribed region from a competitive mapping to both alleles. Diploids show either zero or ⇠ 1*/*2 relative depth over *G2*, while tetraploids show either zero or ⇠ 1*/*4. The x-axis shows ascertained segregating sites from short reads aligned to an assembly containing a single copy of *GFAFL*, with an correction applied for variable sequencing depth. Levels of variation among the *GFAFL* sequences within individuals are much higher for individuals carrying a copy of the *G2* allele. The higher numbers of segregating sites in tetraploids vs. diploids can be explained by the sampling of four vs. two total copies of *GFAFL*. **c)** Expression of the two *GFAFL* alleles across both pistillate and staminate flower development in protandrous and protogynous morphs of *C. paliurus*. Points show biological replicates and lines connect means across replicates. Data reanalyzed from (Qu *et al*. 2023b).

We next examined segregation pattern of the G2 allele in published resequencing data from a larger sample of both diploids and tetraploids (Qu *et al.* 2023a). Among diploids (N=13), genotypes were either G1/G1 (N=4) or G1/G2 (N=9), with no G2/G2 homozygotes seen despite the high frequency of the G2 allele (P=0.056 under the Hardy-Weinberg expectation, x^2^ = 3.64, d.f.=1). Among tetraploids, individuals appear to possess either zero or at most one copy of G2 for every three copies of G1 (Fig. 4b). This pattern is consistent with segregation of G1 and G2 alleles at a single locus during meiosis in tetraploids.

These two alleles deeply diverged, with ⇠8% divergence in amino acid sequence divergence (^dN^/dS = 0.53). Several substitutions occur in conserved regions, in addition to a change in the location of the start codon, suggesting they could encode proteins with divergent function (Fig. S8). We infer that the alleles diverged from one another 45 million years ago (95 %CI 35-54 Mya), consistent with the idea that long-term balancing selection maintains polymorphism at this locus through disassortative mating. This estimate is again more recent than the divergence of *Cyclocarya* and *Pterocarya*, and the *Cyclocarya* GFAFL alleles do not share either of the two conserved GFAFL coding sequence polymorphisms ancestral to *Pterocarya* (Fig. 3a).

### **Cyclocarya* GFAFL* alleles show specialized expression in male and female flowers

To explore the possible functional role of GFAFL in heterodichogamy in *Cyclocarya*, we next examined expression of the two alleles in a published expression dataset representing a developmental time series of staminate and pistillate flowers of both protandrous and protogynous morphs (Qu *et al.* 2023b). The two alleles show striking patterns of unisexual flower-specific expression (Fig. 4c). In protandrous heterozygotes, G2 is more highly expressed during development of pistillate flowers, whereas G1 is more highly expressed during development of staminate flowers, highlighting sub- or neo-functionalization. Yet overall the expression of both alleles in heterozygotes is strongly correlated. These patterns suggest a model wherein both alleles regulate the developmental rate of flowering structures, but one acts as a ‘fast’ allele (G1), while the other acts as a ‘slow’ allele (G2). For this model to be a sufficient basis for heterodichogamy, it would also predict higher expression of the ‘fast’ allele in staminate flowers of protandrous morphs. Consistent with this, we also see that G1 allele is highly upregulated in staminate flowers of protandrous morphs heterozygous for G1 compared to protogynous morphs homozygous for G1, despite its lower relative copy number in heterozygotes. Together, these expression patterns illustrate the potential for heterodichogamy to be controlled by regulatory variation at a single locus with multiple effects on the expression of a single gene. If two alleles differ quantitatively in the rate of promoting the transition to flowering, the combination of (1) sex-specific expression in heterozygotes and (2) morph-specific expression of the recessive allele may be sufficient to generate reciprocal matching of sexes between morphs.

In summary, several lines of evidence support that idea that ancient haplotypes at the GFAFL locus control heterodichogamy in *Cyclocarya*. (1) Two GFAFL alleles segregate at a single locus in association with dichogamy types. (2) Genotype frequencies at this locus are biased in the direction predicted by disassortative mating. (3) These alleles show ancient divergence, consistent with the action of long-term balancing selection. Finally, (4) sex- and morph-specific expression patterns of these GFAFL alleles, together with the gene’s involvement in heterodichogamy in the sister genus, provide strong evidence for a functional role of GFAFL in regulating heterodichogamy. Strikingly, the segregation patterns of these GFAFL alleles reveal a reversed pattern of dominance, where a dominant allele in *Cyclocarya* is associated with protandrous flowering (protandrous heterogamety) in contrast to the protogynous heterogamety seen in *Pterocarya*.

### Convergent genomic structure of the dominant haplotype in **Pterocarya** and **Cyclocarya**

Patterns of allele-specific GFAFL expression in G-locus heterozygotes of both *Pterocarya* and *Cyclocarya* suggest the existence of both cis regulatory divergence between G-locus haplotypes as well as *trans*-acting allele-specific regulation. We noted a striking parallel in haplotype structure between the dominant G-locus haplotypes in both genera that could reflect similar mechanisms of gene regulation.

In *Pterocarya*, pairwise alignments between the dominant and recessive haplotypes in both P. stenoptera and *P. macroptera* revealed the presence of an array of 14-15 ⇠2 kb tandem repeats in both dominant haplotypes (Fig. 5a-c). These tandem repeats do not overlap any predicted genes, but are homologous to sequence present ⇠1.5 kb downstream of GFAFL in both the dominant and recessive haplotypes. In *Cyclocarya*, we detected tandem repeats within the dominant haplotype showing homology to the same location downstream of the transcribed region of GFAFL (Fig. 5d). The repeated sequence occurs in at least three copies in the dominant haplotype of a protandrous tetraploid of *Cyclocarya* (Fig. 5d), and at least two copies in the collapsed assembly from a protandrous diploid (Fig. S13). The repeated regions of *Pterocarya* and *Cyclocarya* show ⇠80% sequence homology to one another over a ⇠1.5 kb region, 700 bp of which aligns to sequence at the syntenic location in *Juglans*.

**Figure 5:**
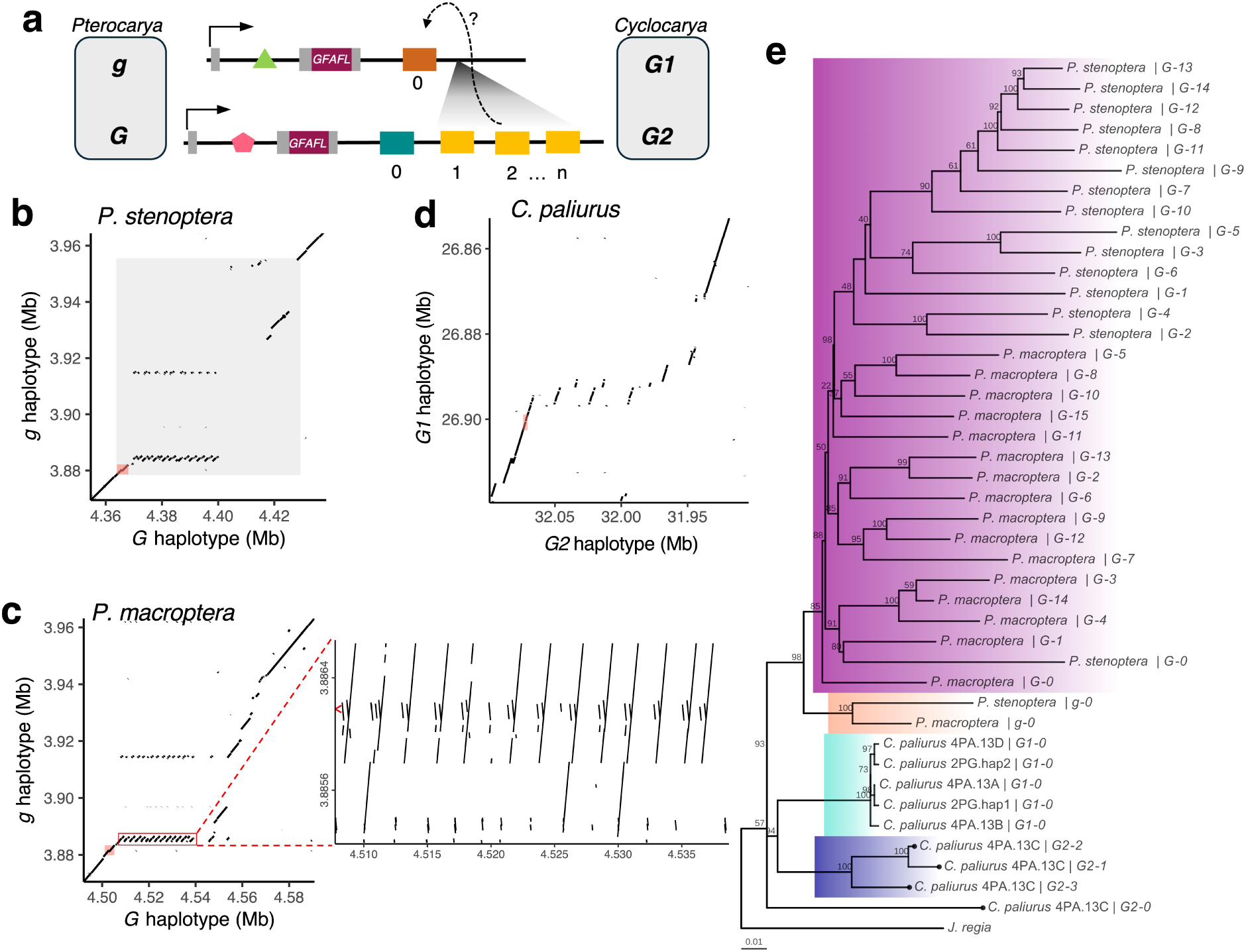
Parallel structure of dominant heterodichogamy haplotypes in **Pterocarya** and **Cyclocarya**. **a)** Simplified schematic of the shared G-locus repeat structure in both genera. Solid arrows indicate the transcription start site for *GFAFL*. Green triangle and pink pentagon represent hypothetical intronic *cis* regulatory elements that differ between haplotypes. Dotted arrow indicates a hypothesized *trans* interaction between the alleles whereby the dominant allele regulates expression of the recessive allele. **b)** Alignment dotplots between the dominant and recessive G-locus of *P. stenoptera*. The gray region indicates the region associated with dichogamy type. Faded red rectangle at bottom left shows the position of *GFAFL* in each assembly. **c)** Alignment dotplot between G-locus haplotypes of *P. macroptera*. Inset at right shows a zoomed in view of the repeats. Red arrow indicates the location of homology in the recessive haplotype to an inverted repeated structure found in both species (see also Fig. S17). **d)** Alignment dotplot between dominant and recessive heterodichogamy haplotypes of *C. paliurus* across the region of segregating copy number variation downstream of *GFAFL* (see Fig. S14). **e)** Maximum likelihood phylogeny of repeat motifs in each assembly rooted with *Juglans* as an outgroup. **Pterocarya** G-locus haplotype sequences are highlighted in purple (*G*) and salmon (*g*). Also highlighted are **Cyclocarya* G2* repeats 1-3 (dark blue) and *G1-0* sequences (light blue). All **Cyclocarya* G2* sequences are indicated with points at tips. Scale bar indicates substitutions per site. Node labels indicate bootstrap support.

We constructed a phylogeny of these repeats and their homologous sequences downstream of GFAFL (Fig. 5e). The dominant haplotype repeats of *Cyclocarya* are reciprocally monophyletic with those found in the dominant haplotype of *Pterocarya*, suggesting these repeats have convergently arisen. The reversed dominance pattern in these genera is also consistent with this interpretation, although an alternative hypothesis is that gene conversion between alternate G-locus haplotypes within each genus has eroded any signal of a common origin of the repeats.

All *Pterocarya* sequences are a well-supported clade (98% bootstrap support), with G haplotype sequences and g haplotype sequences forming two reciprocally monophyletic sister groups (85% and 100% bootstrap support). We found no signal of phylogenetic conservation of specific repeats within the tandem array across these two *Pterocarya* species, indicating considerable turnover in repeat sequence content within species. One or possibly two protogynous individuals of P. stenoptera appear to lack at least part of this repeat array (Fig. S15), potentially reflecting ongoing contraction and expansion. Despite turnover, the strong conservation of this haplotype structure with similar numbers of repeats across the deepest divergence within the genus (and the lack of other notable shared haplotype features) lends support to the idea that these repeats play a role in regulating GFAFL.

In *Cyclocarya*, the allele-specific activation of the G1 copy of GFAFL in staminate flowers in protandrous heterozygotes (Fig. 4c, bottom panels), suggests an interaction in trans between the G2 haplotype and either the transcript of G1 or a *cis* regulatory element on the G1 haplotype. In light of this, it is interesting to note that the *Cyclocarya* G2 repeats are phylogenetically closer to the *Cyclocarya* G1-0 sequence than to the syntenic sequence present on the same haplotype as the repeats themselves (G2-0).

Similarly, in *Pterocarya*, the lack of expression of the g copy of GFAFL in heterozygotes during early development of male flowers, together with its clear expression in gg homozygotes at the same stage, implies that the G haplotype negatively regulates the g copy of GFAFL in heterozgotes. Following a similar logic to above, this suggests a *trans*-acting regulatory mechanism whereby the G haplotype preferentially down-regulates the g copy of GFAFL relative to the G copy. One possible mechanism could involve an RNA intermediary which causes post-transcriptional silencing (Fig. S16), or which suppresses transcription of g through methylation of a *cis* regulatory region. Intriguingly, we noted the presence of a shared ⇠300 bp inverted repeat structure present in multiple copies within the repeat region of both *Pterocarya* dominant haplotypes. These inverted repeats are homologous to the same region in the recessive haplotype of each species and are predicted to form thermodynamically stable hairpin structures (Fig. S17, Axtell and Meyers 2018). This suggests the possibility that allele-specific regulation of GFAFL in G-locus heterozygotes could involve the production of double-stranded small RNAs from the dominant haplotype.

Alternative mechanisms that don’t involve an RNA intermediary are also possible (Matzke *et al.* 2001). For instance, one hypothesis for the expression data in *Pterocarya* is that an enhancer region within the G haplotype outcompetes the g haplotype for recruitment of a limiting shared transcription factor which transiently promotes GFAFL transcription during a specific developmental stage of flowering. Whatever the specific mechanism, these repeated homologous sequences found downstream of GFAFL within dominant haplotypes of both *Pterocarya* and *Cyclocarya* are a promising link for further investigation into the molecular genetic mechanisms of gene regulation that underlie heterodichogamy.

## Discussion

In this work, we documented the occurrence of heterodichogamy *Pterocarya*. Thus, all genera within sub-family Juglandoideae have been confirmed to possess this mating system. Remarkably, our results now demonstrate that among the five genera within Juglandoideae, three distinct genetic loci control heterodichogamy (Groh *et al.* 2025). The genetic basis of heterodichogamy in Platy*Carya* has not yet been identified and may yet involve a separate mechanism. To our knowledge there are no reports of the temporal flowering behavior of subfamily Engelhardiodeae, but considering the similarities of inflorescence structure and pollination mode with Juglandoideae, it seems plausible that heterodichogamy may also be found there also. Neither the *Pterocarya* nor *Cyclocarya* GFAFL alleles, nor the other genetic systems we have previous mapped (Groh *et al.* 2025), are estimated to be ancient enough to be shared across genera. Thus, we cannot definitely rule out convergent evolutionary origins of heterodichogamy in these different genera. However, considering the occurrence of heterodichogamy in all of subfamily Juglandoideae together with its apparent scarcity in related families, it is reasonable to suggest that phenotypic heterodichogamy evolved in the common ancestor of Juglandoideae (Fukuhara and Tokumaru 2014) and has been maintained for >70 million years. If that is the case, the ancestral heterodichogamy mechanism was controlled by a mechanism that was susceptible to subsequent invasion by novel dichogamy-determining alleles. Why might this occur? One possible adaptive explanation could be that heterodichogamy morphs controlled by an ancestral genetic system showed imperfect temporal matching between opposing sexual phases. Such temporal mismatches might arise due to environmental changes that differentially affected male and female flowering time. A novel dichogamy determining allele conferring closer sex matching to the other morph under these conditions could then gain a fitness advantage and takeover control of dichogamy type (Bull 1983). Such alleles may be predisposed to arise at genes that already participate in regulatory networks underlying morph-specific flower development. Consistent with this idea, we see morph-specific temporal expression patterns of previous heterodichogamy candidates in *Juglans* and *Carya* in the flower expression data of *Cyclocarya* (Figs. S18, S19).

As GFAFL regulates heterodichogamy in both *Pterocarya* and *Cyclocarya*, it seems likely that polymorphism at this gene controlled heterodichogamy in their shared ancestor. Phylogenetic evidence suggests that these alleles are reciprocally monophyletic, which would imply both sets of alleles arose after the divergence of these genera (Fig. 4). However, given the ancient age of both sets of alleles and the possibility of re-arrangements between the haplotypes in each genus, we cannot say with confidence whether one of these two systems derived directly from the other, or whether a third GFAFL system was present in their shared ancestor. In either case, a change in the dominance relationships at GFAFL strongly resembles evolutionary changes in the dominance relationships of sex determination systems. A transition from XY to ZW sex determination involving a single gene is found for example in poplars (Müller *et al.* 2020). In that case, the transition appears to be facilitated by different means of regulating the presence/absence of expression of a key regulator of sex-specific development. Allelic turnover in *Pterocarya* and *Cyclocarya* instead appears to involve a transition between two alternate sets of functional protein-coding alleles that differ quantitatively in their regulatory effects on flowering time.

The genetic control of heterodichogamy naturally parallels that of dioecy, but may differ in key ways. While many plant sex chromosomes are homomorphic, heteromorphism seems even less pronounced in heterodichogamy. For one, the apparent lack of phenotypic specialization of dichogamy morphs, other than in their timing, suggests that there may be limited genetic variation with morph-antagonist effects (e.g. protogynous beneficial/ protandrous deleterious alleles or vice versa), precluding an adaptive advantage of recombination suppression that would subsequently trigger degeneration and heteromorphism. Moreover, although disassortative mating renders the dominant homozygote rare, some level of intra-morph mating evidently occurs as viable dominant homozygotes are now known in several genera with mapped heterodichogamy systems. This may enable occasional recombination between dominant haplotypes and a higher rate of purging of deleterious variation compared with permanently-heterozygous heterogametic sex chromosomes.

In contrast to the genetics of dioecy, which involves complete suppression of one sexual function (e.g. via loss-of-function alleles or silencing of a key sex regulator), the molecular basis of heterodichogamy requires quantitative control of gene regulation throughout both male and floral development in order to achieve reciprocal temporal sex matching. How might this be achieved at the genetic level?

One hypothesis is that pleiotropy of a single regulatory change could simultaneously delay flowering of one sex while advancing the other through a resource trade-off (Groh *et al.* 2025). However, if such a trade-off were to exist, then the start of male and female flowering times would likely be negatively correlated within dichogamy types. Our data on the relative developmental rates of male and female inflorescences in *Pterocarya* show the opposite pattern, where male and female developmental rates within each dichogamy type are positively correlated (Fig. 1d). Although the sex-specific phenology variation we see within types is only a subset of the total variation, it is difficult to imagine how the direction of pleiotropy would reverse over some threshold shift in flowering time of either sex.

A second possibility is that variants across multiple genes with sex-specific affects on either male or female flowering time become coupled together on the same haplotype, enabling co-inheritance of variation that delays flowering of one sex while advancing the other. The clear evidence of recombination suppression across multiple genes involved in flowering at the *Carya* G-locus haplotypes lends support to this model.

Our findings in the wingnut group suggest a third possibility. If a single gene regulates the rate of the developmental transition to flowering, and functional alleles vary quantitatively in their effect, heterodichogamy can result from a combination of sex-specific *cis*-regulatory divergence and *trans*-regulation of one allele by the other. The data are clearest in the case of *Cyclocarya*; the G2 allele found in protandrous individuals shows *cis*-activation in female flowers, and the G2 haplotype up-regulates the G1 allele in male flowers. These patterns are consistent with the G2 copy of GFAFL acting a ‘slow’ allele, while the G1 copy acts as a ‘fast’ allele. Our data from *Pterocarya* offer partial support to this model, in that the G haplotype found in protogynous heterozygotes seems to cause allele-specific silencing in trans of the g copy of GFAFL in early male flower development. In this case, the G allele may thus be acting as a ‘slow’ allele compared to a ‘fast’ counterpart, g. It remains possible that GFAFL could play an active role in regulating the timing of female flowering in *Pterocarya*, although we did not detect it here, perhaps due to lack of sampling of the relevant stage. We additionally note that the current data for the *Juglans* locus could also be consistent with this model, but further study is needed.

Our findings show that heterodichogamy can be achieved through diverse genetic pathways, which may facilitate its evolutionary turnover. In wingnuts, and perhaps walnuts, it appears that changes in a single gene may be sufficient to control reciprocal temporal matching of opposing sexual phases. Given this, why then have the current day systems been conserved over vast time periods? Perhaps the molecular architectures of these systems - that is, the putative regulatory repeats - enables sufficient flexibility for adaptation of finely-tuned reciprocal matching, making these systems robust to invasion.

## Author contributions

J.S.G and G.C. conceived of and designed the research. J.S.G. collected phenological data and tissue samples, performed data curation, bioinformatics, formal analysis, and visualization, and wrote the original draft of the manuscript. G.A. isolated nucleic acids from tissue samples. M.W. facilitated tissue collections and contributed phenological observations. G.C. assisted with tissue collections, contributed resources and expertise, and edited the manuscript. All authors reviewed the manuscript and approved submission.

## Data availability

Genome assemblies for P. stenoptera are available at NCBI under accession numbers PRJNA1234227 and PRJNA1234228. We are in the process of uploading the remaining genome assemblies and resequencing data to NCBI and accession numbers will be provided upon resubmission. Scripts used to perform analyses are hosted at https://github.com/jgroh/Pterocarya.

## Acknowledgements

We are grateful to Sonoma Botanical Garden, USDA Wolfskill Experimental Orchard, and Hunter Wilson of UC Davis for providing access to living collections sampled in this study. We thank Pat Brown, Blake Meyers, Chuck Langley, Steven Manchester, Stephen Wright, and members of the Coop lab for helpful comments and discussion. J.S.G. was supported by a research award from the Center for Population Biology, UC Davis.

## Materials and Methods

### Phenotyping

Flower structures were examined in spring of 2024 and dichogamy types were assigned based on the relative developmental stages of pistillate and staminate flowers. Most trees in flower that were observed over multiple time points across the flowering season could be confidently assigned to one dichogamy type. Others were observed on a single day during flowering, and for these dichogamy type was often nonetheless clear but not in all cases. Certainty in the phenotype was greater for trees growing nearby others of the same species experiencing the same local conditions. Binoculars were sometimes required to view flowering structures in the canopy, and here the pink tinge of stigmas was found to be a reliable indicator of the pistillate flowering phase. In concert with an overall assessment of dichogamy type, we took two quantitative measurements for a subset of trees observed. Female catkin lengths of trees in Winters and Davis, CA were measured using a hand ruler within a two-day period in early April near the middle of the flowering season. Male catkin buds on the same set of individuals were measured on Dec. 17 of the same year (during dormancy) under a dissecting microscope, and both length and width were measured using the software ToupView. As a measure of male catkin size, we used each tree’s value along the first principal component from a principal components analysis of both length and width (Fig. 1c).

### Genome Assembly and Annotation

We assembled chromosome-level genomes of one individual each of **Pterocarya* stenoptera, P. fraxinifolia, P. macroptera, and *Cyclocarya* paliurus* (Table S1). The *P. macroptera* and *P. fraxinifolia* assemblies represent the first complete genome assemblies of these species to our knowledge.

Fresh leaf tissue was collected in the field and immediately flash frozen in liquid nitrogen. High Molecular Weight DNA extraction, library preparation, and sequencing on a PacBio Revio machine was performed by MedGenome Inc. (Foster City, CA). For our C. paliurus assembly the DNA was extracted instead from male catkin tissue prior to anther dehiscence. HiFi reads for each individual were assembled into contigs using *hifiasm* 0.19.9 (Cheng *et al*. 2022). As **Cyclocarya** paliurus can be either diploid or polyploid, we first visualized a k-mer count distribution and saw only 2 clear peaks, one at the expected coverage for a haploid assembly given previous estimates of the haploid genome size, and another peak at double this value corresponding to heterozygous regions (additional peaks would be expected for a tetraploid). Thus, we concluded this individual was diploid and proceeded with the same assembly pipeline as for *Pterocarya*. Contig assemblies were then scaffolded using a reference-guided approach using a recent chromosome-level assembly of P. stenoptera which incorporated Hi-C data in scaffolding (Zhang *et al*. 2024d). It is possible that the use of P. stenoptera to scaffold assemblies of *P. fraxinifolia* and *P. macroptera* could lead to assembly errors if there have been large-scale chromosome rearrangements between these species. We expect these to be minimal as chromosome number and synteny over broad chromosomal regions is strongly conserved within Juglandaceae (Lovell *et al*. 2021; Ding *et al*. 2023). For campls, our *P. stenoptera* assembly shows few rearrangements relative to the *J. hindsii* walnut genome (Fig. S5). We assessed the quality of assemblies using continuity measures (e.g. N50) and completeness metrics (e.g. BUSCO). Assembly metrics are reported in Table S1.

We used the BRAKER3 pipeline (Gabriel *et al*. 2024) to annotate our *P. stenoptera* assemblies. The BRAKER3 pipeline incorporates both external evidence from RNA-seq alignments and protein sequences from related species together with *ab initio* predictions. We first identified and soft-masked repetitive sequence in assemblies using the combination of RepeatModeler2 and RepeatMasker (Flynn *et al*. 2020).

We used as input RNA-alignments generated from 12 sequencing libraries derived from male and floral tissue from 10 individuals (this study), combined with 4 published RNA-seq libraries from *P. stenoptera* derived from leaf, stem, root, and fruit (Zhang *et al*. 2024d). We first used STAR (Dobin *et al*. 2013) to perform splice-aware alignment of RNA-seq reads; these alignment files were used as input for BRAKER3. As additional external evidence we used the Viridiplantea partition of the protein OrthoDB v11 protein sequence database (Kuznetsov *et al*. 2023) hosted at https://bioinf.uni-greifswald.de/bioinf/partitioned_odb11/. We note this database includes protein coding sequence from some close relatives, including two species of *Juglans*. To account for the possibility of differential gene content at the locus of interest, we performed annotation separately for our two haplotype resolved assemblies. The annotation pipeline does not predict UTR regions. For the candidate gene *GFAFL*, we manually annotated the UTRs by viewing the coverage extent of RNA-seq reads from our floral tissue libraries. We manually inspected the annotations across haplotypes at the focal locus using gene expression and comparison to other haplotypes in other species. For the annotation shown in Fig. 2, we omit from the visualization one small predicted gene within the G-haplotype which appeared to be an annotation error (see Supplementary Text 2).

We further annotated transposable elements (TEs) in each assembly using the EDTA pipeline (Ou *et al*. 2019). We used bedtools (Quinlan and Hall 2010) to merge overlapping repeat annotations to compute repetitive sequence content in predefined genomic windows. For example in Fig. S7 we compare the repetitive content of G-locus haplotypes to their local genomic context calculated content across 50 flanking windows of equivalent size on either side of each haplotype.

### Species tree inference

We inferred a well-resolved fossil-calibrated phylogeny using whole-genome sequence data for twenty species within Juglandaceae.

### Taxon sampling

Our sample includes five out of six recognized species of *Pterocarya* (according to Plants of the World Online database, https://powo.science.kew.org/, accessed January 2025). A recent study of relationships within *Pterocarya* using reduced representation sequencing of the nuclear genome (Song *et al*. 2020) recovered monophyly of two sections traditionally recognized on the basis of morphology (Manning 1978). Here, we included five *Pterocarya* species for which whole genome sequencing data are available. Three have long-read genome assemblies presented here, P. rhoifolia was shotgun sequenced in Groh *et al*. (2025), and P. hupehensis was shotgun sequenced in Geng *et al*. (2024).

In addition to our sampling of *Pterocarya*, we selected taxa with available whole-genome-sequence data from across Juglandaceae in order capture the phylogenetic diversity within the family. For *Juglans*, we chose species representing four sections traditionally recognized within the genus (Manning 1978). (1) Sect. *Juglans*: J. regia (Marrano *et al*. 2020); (2) sect. Rhysocaryon (black walnuts): *J. hindsii* (Groh *et al*. 2025), J. californica (Fitz-Gibbon *et al*. 2023), J. microcarpa (Zhu *et al*. 2019), and J. nigra (Zhang *et al*. 2021); (3) sect. Cardiocaryon: J. mandhsurica (Li *et al*. 2022); (4) sect. Trachycaryon: J. cinerea (Guzman-Torres *et al*. 2024). For *Carya*, we chose taxa spanning the North American - East Asian divergence (Zhang *et al*. 2013). Species representing the North American clade were pecan (C. illinoinensis) (Lovell *et al*. 2021) and C. ovata (Groh *et al*. 2025), and species representing the East Asian clade were C. sinensis and C. cathayensis (Zhang *et al*. 2024c). Other genomes were from the following sources: **Cyclocarya** paliurus (Qu *et al*. 2023a), Platy*Carya* strobilaceae (Zhou *et al*. 2024); Engelhardia roxburghiana and Rhoiptelea chiliantha (Ding *et al*. 2023).

### Data preparation

The input to our phylogeny inference was a set of orthogroups of protein-coding sequences. For species with published annotations that incorporated RNA-seq data, we used annotations of protein-coding genes directly. For species with no published annotations or having annotations that did not incorporate RNA-seq data from multiple tissues, we used liftoff (Shumate and Salzberg 2021) to identify coding sequence coordinates of protein-coding genes in target assemblies using a reference annotation from a closely related species. For genes with multiple annotated slice variants, we filtered coding sequences so that each gene model was represented by the primary transcript. The final number of transcripts used as input for the orthology search across all species ranged from 27962 to 35370.

For the two species of *Pterocarya* for which we relied on short read resequencing datasets, we constructed pseudo-haploid sequences across annotated exons in our *P. stenoptera* reference genome. To do this, we generate a vcf containing both variant and invariant sites filtered for minimum genotype quality 30 (variant sites only) and a sequencing minimum depth of 5 (all sites). We then used the vcf to extract genotypes within the CDS coordinates of the primary transcripts for all gene models of our *P. stenoptera* annotation using bcftools consensus, encoding sites with zero coverage after filtering (-e ‘FMT/GT==“./.”’ -a “N”-M “N” –mark-del “N”) as ‘N’ in the output files and using IUPAC ambiguity codes to represent heterozygous sites. Consensus sequences were then reverse complemented if the gene annotation was present on the reverse strand in the reference sequence. We then used a custom python script to replace ambiguous IUPAC codes representing heterozygote sites with an appropriate nucleotide chosen at random from the possible bases (except for ‘N’s, which were kept in place). The same procedure was used to construct a pseudohaploid sequence of *Carya* ovata using the C. illinoinensis reference genome.

We then ran OrthoFinder 2.5.5 (Emms and Kelly 2019) with DNA sequence as input (option-d) to identify ortholog groups. 96.0 % of genes were classified into 35,260 orthogroups with a median orthogroup size of 19. OrthoFinder identified 5,089 single-copy orthologs, and 13,652 orthogroups in which all species were represented. The first set of single-copy orthologs was used for the basis of the phylogeny results presented in the main text, and the second set was used as input for a complementary phylogenetic method applied.

### Phylogeny inference

We used both coalescent-based and concatenation approaches to infer the phylogeny. Trees were rooted with *Rhoiptelea chiliantha* as the outgroup. In order to run ASTRAL, for each single copy ortholog group, we performed alignment using muscle (Edgar 2004), and inferred gene trees in IQ-tree (Nguyen *et al*. 2015), specifying a GTR+I+G substitution model and 1000 ultrafast bootstrap replicates. We collapsed nodes with low bootstrap support (< 10%) to polytomies using newick tools https://github.com/xflouris/newick-tools, and then ran ASTRAL-IV v1.19.4.6 (Zhang and Mirarab 2022) on the resulting gene trees to infer a species tree under a model consistent with incomplete lineage sorting. We report Local Posterior Probabilities for internal nodes computed by ASTRAL in the main text. Separately, to perform maximum-likelihood inference, we concatenated alignments of single-copy orthologs to obtain an alignment with 443,387 parsimony-informative sites. The maximum likelihood phylogeny was found in IQ-tree 2.2.5 (Nguyen *et al*. 2015) and in the maintext we report bootstrap confidence values for internal nodes from 1000 replicates of the ultrafast bootstrap implemented in IQ-tree.

Finally, as a complementary approach for phylogenetic inference, we examined the tree inferred by STAG (Emms and Kelly 2018), which is incorporated into the OrthoFinder pipeline. STAG is a gene tree summary method using all ortholog groups in which all species are represented.

All three of these methods inferred the same topology (Fig. 1) with high confidence. The ASTRAL tree had Local Posterior Probabilities of 1 at each internal node, and the maximum likelihood tree had boostrap values of 100% at all internal nodes. Given the strong support for this topology, we treated the topology as fixed in a second step of time-calibration.

### Time-calibration

We performed Bayesian estimation of divergence times using MCMCTree (Yang and Rannala 2006; Reis and Yang 2011). Briefly, this method allows the user to specify priors on node ages, and obtains posterior distributions through Markov-Chain-Monte-Carlo (MCMC) sampling that makes use of an approximation to the likelihood around the maximum likelihood estimate (MLE).

We use fossil information to specify prior distributions on three internal nodes of the Juglandaceae phylogeny (see Supplementary Text 1 for a justification of fossil-based priors and Fig. S3 for visualization). For several nodes we implemented ‘soft’ bounds by placing a small prior mass above and below the bounds for each node, to account for the possibility that nodes are actually older (due to a sparse fossil record) or younger (due to inaccurate fossil placements) than the specified bounds. MCMCTree then uses a birth-death model of speciation to generate priors on the remaining node ages.

We specified a HKY85+r substitution model with 5 discrete rate categories, and an independent-rates relaxed-clock model of substitution rates. The MCMC algorithm was run with a burnin of 50,000 steps, a sampleing frequency of 100 steps, and a sample size of 100,000. We ran two separate chains to check for convergence. We also simulated under the prior (Fig. S3) and compared posterior distributions to prior distributions for nodes of interest (Fig. S4).

We used the fossil-calibrated phylogeny to constrain the date of divergence of the G-locus haplotypes. Given the paucity of shared coding sequence polymorphisms across the *Pterocarya* genus, we chose to use an alignment of the FAFL intron (see Fig. 4). We note the maximum likelihood phylogeny of FAFL coding sequence shows monophyly of the recessive haplotypes of *Pterocarya*, nested within a clade of dominant haplotypes. On the other hand, the maximum likelihood phylogeny of the intron shows reciprocal monophyly of the dominant and recessive haplotypes, which is also consistent with the phylogeny estimate from the G haplotype repeats (Fig. 3). In both cases, **Cyclocarya** is inferred as an outgroup with high bootstrap support when the tree is rooted with additional outgroups.

We used the least-squares dating method implemented IQ-tree (To *et al*. 2016) to obtain point and interval estimates for the divergence time between G and g haplotypes by fixing the divergence of *Pterocarya* and **Cyclocarya** haplotypes from the *Juglans* sequence at the posterior mean and 95% credible interval bounds for the species divergence. The resulting interval estimates for this region for daughter nodes representing within-haplotype divergences in *Pterocarya* all fell within their respective 95% credible intervals for the species divergence times inferred in the genome-wide phylogeny.

Finally, we revisited our date estimates of the divergence ages between haplotypes at both the *Juglans* and *Carya* loci. We used the same molecular clock approaches described in (Groh *et al*. 2025), recalibrated using posterior means and 95% credible intervals for the respective species divergence times that were used in previous calibrations. We obtain estimates of 40.48 (95% CI 33.44-47.12) for the *Juglans* G-locus and 59.47 Mya (95% CI 49.13-69.23) for the *Carya* G-locus, consistent with our previous estimates.

### Short read sequence alignment, variant identification and filtering

For short read data, reads were mapped to the haplotype assemblies of *P. stenoptera* using bwa 0.7.17 (Li and Durbin 2009). Variants were called in bcftools 1.17 (Danecek *et al*. 2021) and genotypes were filtered depending on the purposes of the analysis. SNPs tested for an association with dichogamy type were filtered using criteria of GQ 30 and minimum depth 10. For direct ascertainment of segregating sites in *GFAFL* copies within individuals, we used allele counts directly, setting a frequency cutoff of 0.05 for reads matching the within-individual minor allele.

### Genetic structure and GWAS

We explored genetic structure in our association panel using Principal Components Analysis (PCA) on biallelic SNPs after pruning pairs of sites with genotypic linkage disequilibrium of r^2^ > 0.2. Samples cluster by sampling locality and according to known family relationships (Fig. S2). Both protandrous and protogynous morphs were observed at each sampling locality.

GWAS was performed in GEMMA 0.98.5, which implements a linear mixed effect model to control for relatedness, filtering out sites with MAF< 0.10 and/or with with over 10% of samples having missing data. GWAS was done for alignments of short reads to both haplotypes of *P. stenoptera* assembled with *hifiasm*. Initial results appeared markedly different in that the GWAS against haplotype 2 revealed a single clear association peak, while the GWAS against haplotype 1 revealed many scattered peaks throughout the genome. In the course of our previous work (Groh *et al*. 2025), we observed a similar pattern when performing GWAS against assemblies containing the dominant vs. recessive haplotypes of the pecan G-locus, and determined this was a result of differential transposable element content between the haplotypes at the causal locus. Consistent with this hypothesis, for the GWAS against haplotype 1 where there were multiple peaks, we filtered SNPs that fell within annotations of repetitive sequence and observed that this removed the majority of peaks, while the peak at the syntenic location to that observed against haplotype 2 remained the most consistent with a true association (Fig. S6). A small number of high association SNPs remained at other locations but these lacked a gradual decline in association strength moving outwards from the associated SNP as would be expected based on linkage disequilibrium around a true causal locus embedded within a chromosome. As a further test of the differential TE hypothesis, we extracted local regions around the putative spurious associations in haplotype 1 and used BLAST to detect homology to regions in the haplotype 2 assembly. As expected, those regions we tested showed homology to sequence at the G-locus identified in haplotype 2. All of our subsequent analyses examining the G-locus shown in Fig. 2 and 3 gave us high confidence that this peak is the true causal locus for heterodichogamy in Ptero*Carya*.

### Analysis of polymorphism and divergence within and between haplotypes

We performed whole-genome alignment using anchorwave (Song *et al*. 2022) with settings ‘-R 1 -Q 1’. We calculated nucleotide divergence in 1 kb windows from the anchorwave alignments using a custom R script. In visualizing the divergence in the G-locus region, we filtered out divergence windows containing fewer than 500 aligned bp in or where > 50% of the window was annotated as repeat content in the reference coordinates. We also removed windows that fell within the boundaries of the repeat-rich structural variant after identifying the boundaries using a dotplot from BLAST alignments.

We used minimap2 (Li 2018) to obtain alignments of focal haplotypes in fixed coordinates of one haplotype assembly (i.e. 3a) and called SNPs in bcftools setting ploidy to 1. For SNPs identified between long-read assemblies, we filtered out any singletons in the sample to focus on sites containing information about the topology of the local genealogy. To precisely locate the boundary the boundary of *trans*-species polymorphism, we manually examined SNPs near the border of the region where polymorphism transitions between a species tree topology to one of *trans*-species polymorphism. SNPs that occurred within 3-bp of indels were filtered out.

We extracted a set of *trans*-species polymorphic SNPs between the *P. stenoptera* and *P. macroptera* haplotypes (Fig. 3) and measured measured the fraction of sites at which individuals were heterozygous in a set of independent samples from several Ptero*Carya* species from (Geng *et al*. 2024; Groh *et al*. 2025). As many of these sites were missing in the short read variant calls we considered these fractions as estimates from a larger number of sites and computed confidence intervals using the Agresti-Coull method.

To compare levels of polymorphism within and bewteen G-locus haplotypes, we phased short-read sequencing data using Beagle 5.4 (Browning *et al*. 2021). Phasing was done within species for three species represented by multiple individuals. Phasing results were manually checked for phasing errors across the coding sequence of *GFAFL* by checking for phase switch errors in heterozygotes. We computed pairwise diversity using all pairs of phased haplotypes within each species sample for each haplotype using a custom R script (Fig. 2f).

We computed pairwise dN/dS values bewteen the coding sequence regions of *GFAFL* using the R package ape. We used a custom R script to perform a MacDonald-Kreitman test, where nonsynonymous and synonymous divergence between long-read assemblies of *GFAFL* were compared to the levels of nonsynonymous and synonymous polymorphism seen within g haplotypes after phasing the data. We used polymorphism within g haplotypes as limited polymorphism within G haplotypes restricted the power of the test, nonetheless the result was not significant.

In the above analysis, we omitted a single sample of *P. stenoptera* which showed evidence of chimerism. This sample was derived from a protandrous tree and showed non-zero coverage over the hemizygous region whereas all other protandrous showed effectively zero coverage, but still lower coverage than protogynous samples (Fig. S15b). Outside of the CNV, we directly examined SNPs with the aligned reads for this sample and observed numerous sites where the sample appeared heterozygous for SNPs associated with dichogamy type, but allelic depths at these SNPs deviated from what would be expected for an individual heterozygous for the two G-locus haplotypes. To formally test this, we first ascertained a set of high confidence SNPs where all protogynous Gg heterozygotes were heterozygous. We then measured the average across sites of the relative read depths of the reference vs. alternate alleles within each individual and compared these values among samples. Consistent with our hypothesis that the above protandrous sample was chimeric, we observed it showed a bias in allelic depth distinct from all other samples which showed heterozygosity at these SNPs (see Fig. S20 and associated caption).

To examine segregation of *GFAFL* alleles in *Cyclo*Carya**, we combined our denovo assembly with assembly and resequencing data of C. paliurus from Qu, *et al*. (2023a). We first identified *GFAFL* syntenic regions in these assemblies and plotted average read depth in genomic windows from resequencing samples across across these regions, observing a segregating CNV in both diploids and tetraploids (Figs. S13). We used two complementary approaches to specifically test for segregation of the G1 and G2 alleles of *GFAFL*. First, we aligned all reads of both diploids and tetraploids to a chimeric haploid assembly containing both alleles. In this competitive mapping approach, given the high divergence between the alleles we observed *GFAFL* reads mapped to one or the other alleles with high affinity. For each sample, we took computed the median depths across the transcribed region of these two alleles and measured the relative depth across the alleles to infer copy number in each sample. Second, we aligned all reads to an assembly containing only a single haploid copy of GFAFL to identify segregating sites among GFAFL copies in each sample, in order to test whether samples either contained similar or highly diverged copies of GFAFL. We ascertained a site as segregating if the frequency of the minor allele within each sample was at least 0.05. As these samples varied widely in sequencing depths (< 5 to > 100) and we observed that the number ascertained segregating sites correlated with read depth, we applied a depth correction for the number of segregating sites as follows. The observed number of ascertained segregating sites S*_obs_* is divided by the average across L sites (invariant and variant) of the probability of ascertainment at site ℓ, conditioned on site ℓ segregating and the observed read depth at site ℓ:

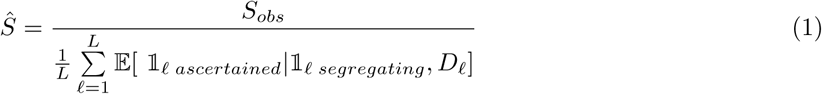

The denominator is the probability of ascertainment for site ℓ conditioned on site ℓ segregating and the observed read depth at site ℓ. This is computed under a binomial sampling model of sequencing reads where p is the within-individual allele frequency (across all copies aligned to the single-copy GFAFL region). For diploids, p = 0.5. For tetraploids, p 2 (0.25, 0.5) and will vary across sites depending on whether a variant exists between homologs or homeologs. For tetraploids with one G2 allele which is highly divergent from G1, there will be only a small proportion of sites where p = 0.5 as most segregating sites will be variants in one of four copies, such that p = 0.25. For tetraploids with zero copies of G2, there will be fewer segregating sites but a greater proportion of these will have p = 0.5. However we do not see a strong signal of divergence between homeologs in the phylogeny and for simplicity we use p = 0.25 for all tetraploids, noting that this choice of p did not qualitatively impact our results.

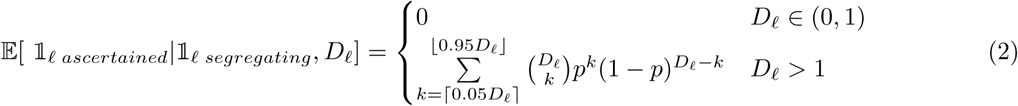

### Relatedness and function of *GFAFL* homologs

We ran the predicted amino acid sequence of GFAFL through InterProScan, which identified a conserved FAF domain (PFAM entry PF11250) in amino acid residues 112-170. The FAF domain is a plant-specific domain of unknown molecular function named for a group of *FANTASTIC FOUR* paralogs in Arabidopsis that were originally identified as regulators of meristem size (Wahl *et al*. 2010). Since then, FAF-like genes have been identified in diverse vascular plants including gymnosperms and lycophytes (Zumajo-Cardona *et al*. 2021). While very few FAF-like genes have been functionally characterized, they have been found to regulate diverse developmental processes including meristem growth (Wahl *et al*. 2010), seed germination, drought tolerance, and primary root growth (Wang *et al*. 2018) and endosperm starch metabolism (Zhang *et al*. 2019).

To examine whether GFAFL showed close homology to any FAF-like genes of known function, we inferred a maximum likelihood phylogeny of FAF-like paralogs. We first identified paralogs within our newly assembled Ptero*Carya* genomes using a protein blast search with the FAF domain of GFAFL as the query sequence. This identified 11 clear paralogs in *P. stenoptera* (blastp E Value<=8e-04). We repeated this paralog search procedure in *Juglans* regia and identified 11 paralogs using the same E value cutoff. We used a similar procedure targeting species in which FAF-like genes have been functionally characterized (Arabidopsis, maize, and tomato) and species selected to represent the diversity of embryophytes (Selaginella and Amborella and Marchantia) in order to provide information about the relatedness of GFAFL to functionally characterized FAF-like genes. We found a single copy FAF-like gene in Marchantia polymorpha, indicating that this gene family was present in the common ancestor of embryophytes (land plants). This sequence was used as an outgroup to root the tree. We performed codon-based alignment of nucleotide sequence encoding the FAF domain (⇠150 nt) using kc-align (https://github.com/davebx/kc-align) and inferred a maximum-likelihood phylogeny in IQtree using a GY+F model of codon evolution and 1000 ultrafast bootstrap replicates (Nguyen *et al*. 2015). We also visualized a pre-constructed gene tree for this gene family in the Panther database (pantherdb.org) which suggested GFAFL is a member of FAF subfamily 17 (SF17, F2E2.18-related).

We then analyzed expression of orthologs of GFAFL in closely related species to provide further information about a conserved biological function. We examined expression data from multiple tissues of the sister genus *Cyclo*Carya** (Zhang *et al*. 2024b). In this dataset which contains vegetative tissues and floral tissue, the GFAFL ortholog is uniquely expressed in flowers (Fig. S9). We note that we later identified expressed GFAFL transcripts in this dataset to originate from both G1 and G2 *Cyclo*Carya** alleles. We then looked in two species from the next most closely related genus, *Juglans*. In *J. regia* we examined expression in 20 tissues using data from (Chakraborty *et al*. 2016), provided by Houston Saxe, UC Davis. The *J. regia* ortholog is predominantly expressed is male catkins. We also examined a dataset from J. nigra (Hardwood Genomics Team 2015) and found a similar pattern of male flower-specific expression (Fig. S10).

We then looked further out to the two closest homologs of GFAFL in Arabidopsis. We note that in the phylogeny shown in Fig. 3 constructed from only the FAF domain, AT1G22110 and AT1G77932 appear equally related to GFAFL. However, alignment over a broader region indicates that AT1G22110 shows much greater sequence similarity. In the expression atlas available at https://www.arabidopsis.org/ showing data from (Klepikova *et al*. 2016), AT1G22110 shows high expression in anthers and weak expression in other tissues. Expression data for this gene from a separate developmental map study (Schmid *et al*. 2005) also show high expression in mature pollen but not other tissues. This suggests that orthologs of GFALF have a conserved functional role in development of staminate flowers and pollen. The other ortholog AT1G77932 shows high expression in flower petals and weak expression in other tissues. Expression data for this gene were not found in the Schmid *et al*. (2005) dataset.

### Expression analysis of *GFAFL*

RNAseq reads were first trimmed using skewer (Jiang *et al*. 2014). We directly aligned RNA-seq reads to the transcriptome using salmon (Patro *et al*. 2017). To study allele-specific expression of GFAFL alleles and mitigate impacts of reference bias, we used a competitive mapping approach by including both GFAFL alleles in the reference transcriptome. To validate this approach, we separately used STAR (Dobin *et al*. 2013) to perform splice-aware alignment of RNAseq reads to both haplotypes and examining SNPs within these reads at sites differing between the two GFAFL alleles.

We first visualized transcript count estimates by tissue and flowering type, and finding that GFAFL was only expressed in male flower tissue proceeded to test for differential expression only for pre-dormancy male catkin buds. We used DESeq2 (Love *et al*. 2014) to obtain transcript counts under a general linear model that corrects for library size and other biases. To test for differential expression of the g allele between gg homozygotes and Gg heterozygotes, we required a separate significance test to account for the difference in copy number. To obtain transcript estimates for a single copy of g we divided the normalized counts for gg homozygotes by two, then fit a generalized linear model to these counts using a negative binomial distribution model and dichogamy type as a predictor. We used a likelihood ratio test from a comparison of this model to one without dichogamy type included as a predictor.

We lacked biological replicates of mature catkin tissue for protandrous morphs of *P. stenoptera*, as this tissue was sampled on a date when the majority of protandrous individuals had already completed pollen shed. Thus, it is possible that overall levels of GFAFL expression may differ between morphs in this tissue at other points during the season.

To test for allele-specific expression in heterozygotes, we used the R function prop.test to test whether the proportion of estimated read counts of each allele differed from 0.5 (two-tailed test). For male bud samples, we ran the test for the two G-locus heterozygotes with sufficient transcript counts for the test to have power. The result was significant in both cases. We report the more conservative P value in the main text, while also noting that the sample with the highest raw transcript count (values shown in the main text apply a library size correction) had 30 estimated transcript counts for GFAFL, all of which were derived from the G allele. In the other two heterozygotes with low transcript counts for the G copy of GFAFL, we still see the same direction of effect, with zero counts of the g copy of GFAFL. The same procedure was done for the mature male catkin tissue, and the result was significant for two out of three heterozygote samples.

In the *Cyclo*Carya** expression data from Qu, *et al*. (2023b), we used salmon to quantify allele-specific transcript counts and DESeq2 to obtain counts counts corrected for variation in library size.

### Analysis of the *GFAFL* region in *Cyclo*Carya**

To investigate the possibility that heterodichogamy in C. paliurus may controlled by variation at the GFAFL locus, we examined short read sequencing depth against the diploid protandrous and protogynous assemblies from Qu, *et al*. (2023a) and our assembly of a diploid protogynous individual. This analysis revealed a segregating copy number variant (CNV) at the location syntenic with the Ptero*Carya* G-locus. Depth of long reads from individuals of known type indicates that this CNV shows an association with dichogamy type (Figs. S13, S14).

We found that the diploid protandrous assembly from Qu, *et al*. (2023a) possessed an annotated copy of GFAFL (CpaM1st14242), and an unannotated copy ca. 100 kb away on the same chromosome. The latter was soft-masked in the original assembly, but we verified it encodes a functional copy of GFAFL using RNAseq (see below). This copy was not found in the assembly of our protogynous individual from the same study, nor in our assembly from a protogynous individual in this study. We determined on the basis of read depth and heterozygosity patterns from short read samples that this haplotype was likely incorrectly assembled and appears to have stitched adjacently together homologous portions of two divergent haplotypes. To further support this, we remapped the long reads from (Qu *et al*. 2023a) to this regions and found discontinuities in coverage between the duplicates, with zero reads spanning a 3-gene cluster that appeared to have a tandem duplication in the assembly (Fig. S13). As this diploid protandrous assembly was generated using a methodology which returns a single collapsed assembly, we concluded that this region was assembled as a chimera from the two divergent haplotypes. We then examined the tetraploid protandrous assembly from Qu *et al*. (2023a), which was generated using a haplotype-specific assembly methodology. We found 4 copies of GFAFL, one of which clustered phylogenetically with the unannotated copy from the diploid protandrous assembly (Figs. 4a, S12). This haplotype in the tetraploid showed no evidence of tandem duplication here, further supporting that GFAFL is single copy in the two haplotypes which are ancestral to the the divergence of diploid and tetraploid lineages.

We next examined in detail four published RNAseq expression datasets including flower tissue from C. paliurus. We found that the samples sequenced in (Chen *et al*. 2019) represent a single time point of the samples reported in the larger time series of (Qu *et al*. 2023a), using PCA to confirm sample identities. Likewise, we noted that the samples sequenced in (Qu *et al*. 2021) appear to be a subset of the data from (Qu *et al*. 2023a), so we did not treat this dataset as independent.

RNAseq data from (Zhang *et al*. 2024d) were not associated with flowering types, and it is not specified whether the sampled tissue originated from male or female flowers. However these data showed clear expression of both GFAFL alleles and confirmed that the expression of GFAFL is flower-specific.

### Analysis of haplotype structure

We used BLAST (Zhang *et al*. 2000) to align G-locus regions of different haplotypes to infer haplotype structure. We extracted the coordinates of repeat regions in G haplotypes and their homologous sequence in g haplotypes from the BLAST alignments, aided by a between-haplotype dotplot showing the alignmed regions. The same procedure was done for the *Cyclo*Carya** haplotypes. These extracted regions were used as BLAST queries to assemblies of *Juglans* and *Carya* to identify orthologous sequence at syntenic regions in other assemblies. As in Ptero*Carya* the sequences identified in *Juglans* and *Carya* are located approximately 1 kb downstream of the 3’ UTR of the GFAFL orthologs. We used muscle (Edgar 2004) to align these sequences and then manually curated these alignments to remove aligned regions where homology was dubious, namely in regions with indels and at the edges of the alignments. A maximum likelihood phylogeny was inferred from this alignment using IQ-tree (Nguyen *et al*. 2015) with 1000 ultrafast bootstrap replicates.

Given the apparent *trans*-acting allele-specific regulation associated with the presence of the G haplotype repeats in Ptero*Carya* and the G2 repeats in *Cyclo*Carya**, we considered the possibility of post-transcriptional silencing, and so tested for possible sequence homology over at least 20 bp between the repeated regions in the dominant haplotypes of both Ptero*Carya* and *Cyclo*Carya** and the transcribed regions of GFAFL on both haplotypes using a blastn-short search with lenient alignment parameters to detect homology between short sequences. In Cylo*Carya*, we found no compelling matches between the entire hemizygous region and the transcribed region of either GFAFL allele.

In *P. stenoptera*, we identified no regions that showed sequence matching between any part of the transcribed region of the dominant haplotype copy of GFAFL and the entirety of the G-repeat region. However, we found that several G-repeats that showed exact sequence matching over 25-30 bp to a site within the 5’ UTR intron of only the recessive copy of GFAFL, roughly 770 bp upstream of the start codon (Fig. S16). Interestingly, at the the same local location within the GFAFL intron in *P. macroptera*, we found several G-repeats matching the recessive but not dominant FAFL intron over 20-21 bp. This sequence motif is not unique to this genomic region, and we considered the possibility that it could represent a regulatory motif throughout the genome. Using a blastn-short search against the *P. stenoptera* genome with the 25 bp intron-matching motif common to three *P. stenoptera* G repeats, we identified 5099 copies of this sequence using an E value cutoff of 0.0015 (corresponding to perfect sequence matching over at least 21 bp or one mismatch over 25 bp). We then intersected these with 300 bp and 2 kb regions flanking all coding sequence regions, as well as with coding sequence itself and introns within coding sequence. We found a significant enrichment (P < 2.2e-16) of the motif within both upstream and downstream 2 kb flanking regions (but not 300 bp flanking regions), suggesting this could be a regulatory motif. To test for significance of this pattern, we asked whether the proportion of motifs identified within the genome overlapped a given feature more than expected given the proportion of the genome constituted by that feature. However, this motif is A-T rich, which may suggest it does not encode a stable small RNA. Also, despite its high copy number in genomes of two *Juglans* species, we detected few or no sequences matching the motif in small RNA sequencing libraries derived from the homologous tissue (Groh *et al*. 2025).

To investigate the possibility of small RNAs deriving from the G haplotype, we used alignment dotplots to look for inverted repeat structures. We identified an inverted repeat at the same location in both Ptero*Carya* species (Fig. S17). We used the RNAfold web server to predict secondary structures of this inverted repeat.

## Notes

### Competing Interest Statement

The authors have declared no competing interest.

## References

Akagi T, Fujita N, Shirasawa K, Tanaka H, Nagaki K, Masuda K, Horiuchi A, Kuwada E, Kawai K, Kunou R et al. 2025. Rapid and dynamic evolution of a giant Y chromosome in Silene latifolia. Science. 387:637–643.

APG IV. 2016. An update of the Angiosperm Phylogeny Group classification for the orders and families of flowering plants: APG IV. Botanical journal of the Linnean Society. 181:1–20.

Axtell MJ, Meyers BC. 2018. Revisiting criteria for plant microRNA annotation in the era of big data. The Plant Cell. 30:272–284.

Bachtrog D. 2013. Y-chromosome evolution: Emerging insights into processes of Y-chromosome degeneration. Nature Reviews Genetics. 14:113–124.

Bai WN, Zeng YF, Zhang DY. 2007. Mating patterns and pollen dispersal in a heterodichogamous tree, *Juglans* mandshurica (Juglandaceae). New Phytologist. 176:699–707.

Belt ES, Hartman JH, Diemer JA, Kroeger TJ, Tibert NE, Curran HA. 2004. Unconformities and age relationships, Tongue River and older members of the Fort Union Formation (Paleocene), western Williston Basin, USA. Rocky Mountain Geology. 39:113–140.

Browning BL, Tian X, Zhou Y, Browning SR. 2021. Fast two-stage phasing of large-scale sequence data. The American Journal of Human Genetics. 108:1880–1890.

Brunet J, Charlesworth D. 1995. Floral sex allocation in sequentially blooming plants. Evolution. 49:70–79. Bull JJ. 1983. Evolution of sex determining mechanisms.. The Benjamin/Cummings Publishing Company, Inc.

Chakraborty S, Britton M, Martínez-Garćıa P, Dandekar AM. 2016. Deep RNA-Seq profile reveals biodiversity, plant–microbe interactions and a large family of NBS-LRR resistance genes in walnut (*Juglans* regia) tissues. AMB Express. 6:1–13.

Chen X, Mao X, Huang P, Fang S. 2019. Morphological characterization of flower buds development and related gene expression profiling at bud break stage in heterodichogamous *Cyclocarya* paliurus (Batal.) lljinskaja. Genes. 10:818.

Cheng H, Jarvis ED, Fedrigo O, Koepfli KP, Urban L, Gemmell NJ, Li H. 2022. Haplotype-resolved assembly of diploid genomes without parental data. Nature Biotechnology. 40:1332–1335.

Danecek P, Bonfield JK, Liddle J, Marshall J, Ohan V, Pollard MO, Whitwham A, Keane T, McCarthy SA, Davies RM et al. 2021. Twelve years of SAMtools and BCFtools. GigaScience. 10:giab008.

Darwin C. 1877. The Different Forms of Flowers on Plants of the Same Species. John Murray.

Delpino F. 1874. Ulteriori osservazioni e considerazioni sulla dicogamia nel regno vegetale. Appendice. Dimorfismo nel noce (Juglans regia) e pleiontismo nelle piante. Atti della Societa Italiana Scientia Naturale. 17:402–407.

Ding YM, Pang XX, Cao Y, Zhang WP, Renner SS, Zhang DY, Bai WN. 2023. Genome structure-based Juglandaceae phylogenies contradict alignment-based phylogenies and substitution rates vary with DNA repair genes. Nature Communications. 14:617.

Dobin A, Davis CA, Schlesinger F, Drenkow J, Zaleski C, Jha S, Batut P, Chaisson M, Gingeras TR. 2013. STAR: ultrafast universal RNA-seq aligner. Bioinformatics. 29:15–21.

Edgar RC. 2004. MUSCLE: multiple sequence alignment with high accuracy and high throughput. Nucleic Acids Research. 32:1792–1797.

Emms D, Kelly S. 2018.STAG: species tree inference from all genes. BioRxiv. p. 267914.

Emms DM, Kelly S. 2019. OrthoFinder: phylogenetic orthology inference for comparative genomics. Genome Biology. 20:1–14.

Endress PK. 2020. Structural and temporal modes of heterodichogamy and similar patterns across angiosperms. Botanical Journal of the Linnean Society. 193:5–18.

Fisher RA. 1930. The genetical theory of natural selection. Clarendon Press.

Fitz-Gibbon S, Mead A, O’Donnell S, Li ZZ, Escalona M, Beraut E, Sacco S, Marimuthu MP, Nguyen O, Sork VL. 2023. Reference genome of California walnut, Juglans californica, and resemblance with other genomes in the order Fagales. Journal of Heredity. esad036.

Flynn JM, Hubley R, Goubert C, Rosen J, Clark AG, Feschotte C, Smit AF. 2020. RepeatModeler2 for automated genomic discovery of transposable element families. Proceedings of the National Academy of Sciences. 117:9451–9457.

Franks SJ, Sim S, Weis AE. 2007. Rapid evolution of flowering time by an annual plant in response to a climate fluctuation. Proceedings of the National Academy of Sciences. 104:1278–1282.

Friis EM, Pedersen KR, Schönenberger J. 2006. Normapolles plants: a prominent component of the Cretaceous rosid diversification. Plant Systematics and Evolution. 260:107–140.

Fukuhara T, Tokumaru Si. 2014. Inflorescence dimorphism, heterodichogamy and thrips pollination in Platy*Carya* strobilacea (Juglandaceae). Annals of Botany. 113:467–476.

Gabriel L, Brna T, Hoff KJ, Ebel M, Lomsadze A, Borodovsky M, Stanke M. 2024. BRAKER3: Fully automated genome annotation using RNA-seq and protein evidence with GeneMark-ETP, AUGUSTUS, and TSEBRA. Genome Research. 34:769–777.

Ganders FR. 1979. The biology of heterostyly. New Zealand Journal of Botany. 17:607–635.

Gaudinier A, Blackman BK. 2020. Evolutionary processes from the perspective of flowering time diversity. New Phytologist. 225:1883–1898.

Geng F, Zhang X, Ma J, Liu H, Ye H, Hao F, Liu M, Dang M, Zhou H, Li M et al. 2024. Genome assembly and winged fruit gene regulation of Chinese wingnut: insights from genomic and transcriptomic analyses. Genomics, Proteomics & Bioinformatics. qzae087.

Gleeson SK. 1982. Heterodichogamy in walnuts: inheritance and stable ratios. Evolution. pp. 892–902.

Groh JS, Vik DC, Davis M, Monroe JG, Stevens KA, Brown PJ, Langley CH, Coop G. 2025. Ancient structural variants control sex-specific flowering time morphs in walnuts and hickories. Science. 387:eado5578.

Guzman-Torres CR, Trybulec E, LeVasseur H, Akella H, Amee M, Strickland E, Pauloski N, Williams M, Romero-Severson J, Hoban S et al. 2024. Conserving a threatened North American walnut: a chromosome-scale reference genome for butternut (*Juglans* cinerea). G3: Genes, Genomes, Genetics. 14:jkad189.

Hall MC, Willis JH. 2006. Divergent selection on flowering time contributes to local adaptation in Mimulus guttatus populations. Evolution. 60:2466–2477.

Hardwood Genomics Team. 2015. Comparative genomics of environmental stress responses in North American hardwoods. https://www.ncbi.nlm.nih.gov/bioproject/273267.

Hěrmanová Z, Kvček J, Friis EM. 2011. Budvaricarpus serialis Knobloch & Mai, an unusual new member of the Normapolles complex from the Late Cretaceous of the Czech Republic. International Journal of Plant Sciences. 172:285–293.

Jiang H, Lei R, Ding SW, Zhu S. 2014. Skewer: a fast and accurate adapter trimmer for next-generation sequencing paired-end reads. BMC Bioinformatics. 15:1–12.

Klepikova AV, Kasianov AS, Gerasimov ES, Logacheva MD, Penin AA. 2016. A high resolution map of the Arabidopsis thaliana developmental transcriptome based on rna-seq profiling. The Plant Journal. 88:1058– 1070.

Kuznetsov D, Tegenfeldt F, Manni M, Seppey M, Berkeley M, Kriventseva EV, Zdobnov EM. 2023. OrthoDB v11: annotation of orthologs in the widest sampling of organismal diversity. Nucleic Acids Research. 51:D445–D451.

Leigh EGJ. 1976. Sex ratio, sex change, and natural selection. Proceedings of the National Academy of Sciences. 73:3656–3660.

Li H. 2018. Minimap2: pairwise alignment for nucleotide sequences. Bioinformatics. 34:3094–3100.

Li H, Durbin R. 2009. Fast and accurate short read alignment with Burrows–Wheeler transform. Bioinformatics. 25:1754–1760.

Li X, Cai K, Zhang Q, Pei X, Chen S, Jiang L, Han Z, Zhao M, Li Y, Zhang X et al. 2022. The Manchurian walnut genome: insights into juglone and lipid biosynthesis. GigaScience. 11:giac057.

Lim CW, Bae Y, Lee SC. 2022. Differential role of Capsicum annuum *FANTASTIC FOUR*-like gene CaFAF1 on drought and salt stress responses. Environmental and Experimental Botany. 199:104887.

Lloyd DG, Webb C. 1986. The avoidance of interference between the presentation of pollen and stigmas in angiosperms I. Dichogamy. New Zealand Journal of Botany. 24:135–162.

Love M, Anders S, Huber W et al. 2014. Differential analysis of count data–the DESeq2 package. Genome Biol. 15:10–1186.

Lovell JT, Bentley NB, Bhattarai G, Jenkins JW, Sreedasyam A, Alarcon Y, Bock C, Boston LB, Carlson J, Cervantes K et al. 2021. Four chromosome scale genomes and a pan-genome annotation to accelerate pecan tree breeding. Nature Communications. 12:4125.

Luza JG, Polito VS. 1988. Microsporogenesis and anther differentiation in *Juglans* regia L.: A developmental basis for heterodichogamy in walnut. Botanical Gazette. 149:30–36.

Lyson TR, Miller I, Bercovici A, Weissenburger K, Fuentes A, Clyde W, Hagadorn J, Butrim M, Johnson K, Fleming R et al. 2019. Exceptional continental record of biotic recovery after the Cretaceous–Paleogene mass extinction. Science. 366:977–983.

Manchester SR. 1987. The fossil history of the Juglandaceae. Monographs in Systematic Botany from the Missouri Botanical Garden. 21:1–137.

Manchester SR. 1991. Cruciptera, a new juglandaceous winged fruit from the Eocene and Oligocene of western North America. Systematic Botany. pp. 715–725.

Manchester SR, Dilcher DL. 1982. Pterocaryoid fruits (Juglandaceae) in the Paleogene of North America and their evolutionary and biogeographic significance. American Journal of Botany. 69:275–286.

Manchester SR, Dilcher DL. 1997. Reproductive and vegetative morphology of Polyptera (Juglandaceae) from the Paleocene of Wyoming and Montana. American Journal of Botany. 84:649–663.

Manning WE. 1978. The classification within the Juglandaceae. Annals of the Missouri botanical garden. pp. 1058–1087.

Manos PS, Soltis PS, Soltis DE, Manchester SR, Oh SH, Bell CD, Dilcher DL, Stone DE. 2007. Phylogeny of extant and fossil Juglandaceae inferred from the integration of molecular and morphological data sets. Systematic Biology. 56:412–430.

Mao X, Liu J, Li X, Qin J, Fu X. 2016. Flowering biological characteristics and mating system in immature plantations of heterodichogamous *Cyclocarya* paliurus. Journal of Nanjing Forestry University. 59:47.

Marrano A, Britton M, Zaini PA, Zimin AV, Workman RE, Puiu D, Bianco L, Pierro EAD, Allen BJ, Chakraborty S et al. 2020. High-quality chromosome-scale assembly of the walnut (*Juglans* regia L.) reference genome. GigaScience. 9:giaa050.

Matzke M, Mette MF, Jakowitsch J, Kanno T, Moscone EA, Van Der Winden J, Matzke AJ. 2001. A test for transvection in plants: DNA pairing may lead to *trans*-activation or silencing of complex heteroalleles in tobacco. Genetics. 158:451–461.

Moraga C, Branco C, Rougemont Q, Jedlčka P, Mendoza-Galindo E, Veltsos P, Hanique M, Rodŕıguez de la Vega RC, Tannier E, Liu X et al. 2025. The Silene latifolia genome and its giant Y chromosome. Science. 387:630–636.

Mu XY, Tong L, Sun M, Zhu YX, Wen J, Lin QW, Liu B. 2020. Phylogeny and divergence time estimation of the walnut family (Juglandaceae) based on nuclear RAD-Seq and chloroplast genome data. Molecular Phylogenetics and Evolution. 147:106802.

Müller NA, Kersten B, Leite Montalvāo AP, Mähler N, Bernhardsson C, Bräutigam K, Carracedo Lorenzo Z, Hoenicka H, Kumar V, Mader M, et al. 2020. A single gene underlies the dynamic evolution of poplar sex determination. Nature Plants. 6:630–637.

Munday PL, Buston PM, Warner RR. 2006. Diversity and flexibility of sex-change strategies in animals. Trends in Ecology & Evolution. 21:89–95.

Nguyen LT, Schmidt HA, Von Haeseler A, Minh BQ. 2015. IQ-TREE: a fast and effective stochastic algorithm for estimating maximum-likelihood phylogenies. Molecular Biology and Evolution. 32:268–274.

Oginuma K. 1999. Karyomorphology and evolution in Juglandales: a Review. Acta Phytotax. Geobot.. 50:229–241.

Ou S, Su W, Liao Y, Chougule K, Agda JR, Hellinga AJ, Lugo CSB, Elliott TA, Ware D, Peterson T et al. 2019. Benchmarking transposable element annotation methods for creation of a streamlined, comprehensive pipeline. Genome Biology. 20:1–18.

Pannell JR, Verdú M. 2006. The evolution of gender specialization from dimorphic hermaphroditism: paths from heterodichogamy to gynodioecy and androdioecy. Evolution. 60:660–673.

Patro R, Duggal G, Love MI, Irizarry RA, Kingsford C. 2017. Salmon provides fast and bias-aware quantification of transcript expression. Nature Methods. 14:417–419.

Pendleton RL, Freeman DC, McArthur ED, Sanderson SC. 2000. Gender specialization in heterodichogamous Grayia brandegei (Chenopodiaceae): evidence for an alternative pathway to dioecy. American Journal of Botany. 87:508–516.

Pigg KB, DeVore ML, Taylor W. 2023. New features of *Cyclocarya* brownii Manchester & Dilcher from the Late Paleocene of North Dakota, USA. International Journal of Plant Sciences. 184:282–303.

Puttick MN. 2019. MCMCtreeR: functions to prepare MCMCtree analyses and visualize posterior ages on trees. Bioinformatics. 35:5321–5322.

Qu Y, Kong W, Wang Q, Fu X. 2021. Genome-wide identification MIKC-Type MADS-Box gene family and their roles during development of floral buds in wheel wingnut (*Cyclocarya* paliurus). International Journal of Molecular Sciences. 22:10128.

Qu Y, Shang X, Fang S, Zhang X, Fu X. 2023a. Genome assembly of two diploid and one auto-tetraploid *Cyclocarya* paliurus genomes. Scientific Data. 10:507.

Qu Y, Shang X, Zeng Z, Yu Y, Bian G, Wang W, Liu L, Tian L, Zhang S, Wang Q et al. 2023b. Whole-genome duplication reshaped adaptive evolution in a relict plant species, *Cyclocarya* paliurus. Genomics, Proteomics & Bioinformatics. 21:455–469.

Quinlan AR, Hall IM. 2010. BEDTools: a flexible suite of utilities for comparing genomic features. Bioinformatics. 26:841–842.

Reis Md, Yang Z. 2011. Approximate likelihood calculation on a phylogeny for Bayesian estimation of divergence times. Molecular biology and evolution. 28:2161–2172.

Renner SS. 2001. How common is heterodichogamy? Trends in Ecology & Evolution. 16:595–597.

Rhein HS, Sreedasyam A, Cooke P, Velasco-Cruz C, Grimwood J, Schmutz J, Jenkins J, Kumar S, Song M, Heerema RJ et al. 2023. Comparative transcriptome analyses reveal insights into catkin bloom patterns in pecan protogynous and protandrous cultivars. PloS One. 18:e0281805.

Routley MB, Bertin RI, Husband BC. 2004. Correlated evolution of dichogamy and self-incompatibility: a phylogenetic perspective. International Journal of Plant Sciences. 165:983–993.

Schmid M, Davison TS, Henz SR, Pape UJ, Demar M, Vingron M, Schölkopf B, Weigel D, Lohmann JU. 2005. A gene expression map of Arabidopsis thaliana development. Nature Genetics. 37:501–506.

Shumate A, Salzberg SL. 2021. Liftoff: accurate mapping of gene annotations. Bioinformatics. 37:1639–1643.

Song B, Marco-Sola S, Moreto M, Johnson L, Buckler ES, Stitzer MC. 2022. AnchorWave: Sensitive alignment of genomes with high sequence diversity, extensive structural polymorphism, and whole-genome duplication. Proceedings of the National Academy of Sciences. 119:e2113075119.

Song YG, Li Y, Meng HH, Fragnìere Y, Ge BJ, Sakio H, Yousefzadeh H, Bétrisey S, Kozlowski G. 2020. Phylogeny, taxonomy, and biogeography of *Pterocarya* (Juglandaceae). Plants. 9:1524.

Song YG, Walas L, Pietras M, Ŝam HV, Yousefzadeh H, Ok T, Farzaliyev V, Worobiec G, Worobiec E, Stachowicz-Rybka R, et al. 2021. Past, present and future suitable areas for the relict tree *Pterocarya* fraxinifolia (Juglandaceae): Integrating fossil records, niche modeling, and phylogeography for conservation. European Journal of Forest Research. 140:1323–1339.

Stolle E, Pracana R, Howard P, Paris CI, Brown SJ, Castillo-Carrillo C, Rossiter SJ, Wurm Y. 2019. Degenerative expansion of a young supergene. Molecular Biology and Evolution. 36:553–561.

Stults DZ, Tiffney BH, Axsmith BJ. 2022. New observations on the last *Pterocarya* (Juglandaceae) occurrences in eastern North America. International Journal of Plant Sciences. 183:380–392.

Sutton J. 2019. ‘Cyclocarya paliurus’ from the website Trees and Shrubs Online. https://www.treesandshrubsonline.org/articles/Cyclocarya/ International Dendrology Society.

Thompson T, Romberg L. 1985. Inheritance of heterodichogamy in pecan. Journal of Heredity. 76:456–458.

To TH, Jung M, Lycett S, Gascuel O. 2016. Fast dating using least-squares criteria and algorithms. Systematic biology. 65:82–97.

Wahl V, Brand LH, Guo YL, Schmid M. 2010. The *FANTASTIC FOUR* proteins influence shoot meristem size in Arabidopsis thaliana. BMC Plant Biology. 10:1–12.

Wang K, He J, Zhao Y, Wu T, Zhou X, Ding Y, Kong L, Wang X, Wang Y, Li J et al. 2018. Ear1 negatively regulates aba signaling by enhancing 2c protein phosphatase activity. The Plant Cell. 30:815–834.

Warner RR. 1975. The adaptive significance of sequential hermaphroditism in animals. The American Naturalist. 109:61–82.

Wolfe JA. 1973. Fossil forms of Amentiferae. Brittonia. 25:334–355.

Yan H, Zhou P, Wang W, Ye JF, Tan SL, Guo CC, Zhang WG, Zhu ZW, Liu YZ, Xiang XG. 2024. Biogeographic history of *Pterocarya* (Juglandaceae) inferred from phylogenomic and fossil data. Journal of Systematics and Evolution. 62:1165–1176.

Yang Z, Rannala B. 2006. Bayesian estimation of species divergence times under a molecular clock using multiple fossil calibrations with soft bounds. Molecular biology and evolution. 23:212–226.

Yu RM, Zhang N, Zhang BW, Liang Y, Pang XX, Cao L, Chen YD, Zhang WP, Yang Y, Zhang DY et al. 2023. Genomic insights into biased allele loss and increased gene numbers after genome duplication in autotetraploid *Cyclocarya* paliurus. BMC biology. 21:168.

Zhang C, Mirarab S. 2022. Weighting by gene tree uncertainty improves accuracy of quartet-based species trees. Molecular Biology and Evolution. 39:msac215.

Zhang C, Rabiee M, Sayyari E, Mirarab S. 2018. ASTRAL-III: polynomial time species tree reconstruction from partially resolved gene trees. BMC bioinformatics. 19:15–30.

Zhang D, Ai G, Ji K, Huang R, Chen C, Yang Z, Wang J, Cui L, Li G, Tahira M et al. 2024a. EARLY FLOWERING is a dominant gain-of-function allele of *FANTASTIC FOUR* 1/2c that promotes early flowering in tomato. Plant Biotechnology Journal. 22:698–711.

Zhang JB, Li RQ, Xiang XG, Manchester SR, Lin L, Wang W, Wen J, Chen ZD. 2013. Integrated fossil and molecular data reveal the biogeographic diversification of the eastern Asian-eastern North American disjunct hickory genus (*Carya* Nutt.). PLoS One. 8:e70449.

Zhang Sy, Peng Yq, Xiang Gs, Song Wl, Feng L, Jiang Xy, Li Xj, He Sm, Yang Sc, Zhao Y et al. 2024b. Functional characterization of genes related to triterpene and flavonoid biosynthesis in *Cyclocarya* paliurus. Planta. 259:50.

Zhang WP, Cao L, Lin XR, Ding YM, Liang Y, Zhang DY, Pang EL, Renner SS, Bai WN. 2021. Dead-end hybridization in walnut trees revealed by large-scale genomic sequence data. Molecular Biology and Evolution. 39:msab308.

Zhang WP, Ding YM, Cao Y, Li P, Yang Y, Pang XX, Bai WN, Zhang DY. 2024c. Uncovering ghost introgression through genomic analysis of a distinct eastern asian hickory species. The Plant Journal. 119:1386–1399.

Zhang X, Mogel KJHv, Lor VS, Hirsch CN, De Vries B, Kaeppler HF, Tracy WF, Kaeppler SM. 2019. Maize sugary enhancer1 (se1) is a gene affecting endosperm starch metabolism. Proceedings of the National Academy of Sciences. 116:20776–20785.

Zhang Z, Schwartz S, Wagner L, Miller W. 2000. A greedy algorithm for aligning DNA sequences. Journal of Computational biology. 7:203–214.

Zhang ZY, Xia HX, Yuan MJ, Gao F, Bao WH, Jin L, Li M, Li Y. 2024d. Multi-omics analyses provide insights into the evolutionary history and the synthesis of medicinal components of the Chinese wingnut. Plant Diversity. 46:309–320.

Zhao JL, Dong Y, Huang ADAD, Duan SC, Peng XC, Liao H, Chen J, Luo YL, Lan QY, Wang YL et al. 2025. Ginger genome enables identification of SMPED1 causing sex-phase synchrony and outcrossing in a flowering plant. Preprint (Version 1, 13 Feb 2025). Research Square. 10.21203/rs.3.rs-5849960/v1.

Zhou H, Hu Y, Ebrahimi A, Liu P, Woeste K, Zhao P, Zhang S. 2021. Whole genome based insights into the phylogeny and evolution of the Juglandaceae. BMC Ecology and Evolution. 21:1–16.

Zhou H, Zhang X, Liu H, Ma j, Hao F, Ye H, Wang Y, Zhang S, Yue M, Zhao P. 2024. Chromosome-level genome assembly of Platy*Carya* strobilacea. Scientific Data. 11:269.

Zhu T, Wang L, You FM, Rodriguez JC, Deal KR, Chen L, Li J, Chakraborty S, Balan B, Jiang CZ et al. 2019. Sequencing a *Juglans* regia ⇥ J. microcarpa hybrid yields high-quality genome assemblies of parental species. Horticulture Research. 6:55.

Zumajo-Cardona C, Little DP, Stevenson D, Ambrose BA. 2021. Expression analyses in Ginkgo biloba provide new insights into the evolution and development of the seed. Scientific Reports. 11:21995.

