## Supplementary Material for "Allelic turnover and dominance reversal at a single-gene balanced polymorphism controlling heterodichogamous flowering in wingnuts (Juglandaceae)"

#### Supplementary Text 1

We use fossil information to specify prior distributions on internal nodes of the Juglandaceae phylogeny. In each case, distributions were specifically tailored reflect our uncertainty in both fossil age and assignment. We also implemented MCMCTree's ability to use 'soft' bounds by placing a small prior mass below the minimum and/or above the maximum bounds, accounting for the possibility that nodes are actually younger (i.e. due to inaccurate fossil assignment) or older (i.e. due to a sparse preservation/detection of fossils) than the specified bounds. MCMCTree then uses a birth-death model to generate a joint prior on all internal node ages lacking manually specified priors. Seven fossil-calibrated nodes are listed below in order from oldest to youngest.

We note that in general, specifying a minimum age constraint for a daughter node implies the same minimum age constraint for a parent node. In this sense, a fossil constraining the minimum age of the parent node, which is younger than or equal in age to a fossil constraining the daughter node, would provide no additional information. However, if the minimum bound on a daughter node is 'soft', this relaxes the constraint on parent nodes, and a younger fossil constraining the parent node with a 'hard' minimum bound can provide non-redundant information. Following this logic, here we use minimum age constraints on both the Juglandoideae-Engelhardiodeae divergence (parent node) and the *Pterocarya*-*Cyclocarya* divergence (daughter node) based on fossils of similar age, but treat the minimum constraint on the former as 'hard' and the minimum age constraint on the latter as 'soft' (justification provided below). We used the R package MCMCTreeR (Puttick 2019) to identify parameters specifying the desired prior distributions. The joint prior on all internal nodes is shown in Fig. S3, with a subset of nodes numbered in correspondence with the list of calibration points below.

1. Crown age of Juglandaceae
   Min: 70 Mya
   Max: 96.5 Mya

   **Justification:**
   Here, we follow the Angiosperm Phylogeny Group IV classification of Juglandaceae which includes *Rhoiptelea* (APG IV 2016). Elsewhere, *Rhoiptelea* is regarded as the sole member of the family Rhoipteleaceae; unlike other Juglandaceae it possesses cosexual flowers and a superior ovary.

   Juglandaceae is considered to have likely derived from members of the Normapolles complex, a diverse fossil assemblage of the Late Cretaceous and Tertiary with morphological affinities to the modern order Fagales (Friis *et al.* 2006). Normapolles pollen and associated flower structures first appear in the Upper Cenomanian and are abundant throughout the remainder of the Cretaceous (Wolfe 1973; Friis *et al.* 2006). Considering Fagales (which includes 6 other families in addition to Juglandaceae) to have derived from the members of the Normapolles complex, pollen of which is common in late Cretaceous sediments, we set the Upper Cenomanian as a maximum age for this node, as this represents the earliest occurrence of Normapolles fossils.

   Fossil flowers of *Budvaricarpus* from the Late Turonian-Santonian (minimum age 83.6 Mya) as described by (Heřmanová *et al.* 2011) bear close resemblance to those of extant *Rhoiptelea*. These authors suggested these flowers and those of *Caryanthus*, an affiliated member of the Normapolles group, as possible basal lineages of Juglandaceae (here, including *Rhoiptelea*). Other workers have noted close affinity of pollen from the Late Cretaceous to that of *Rhoiptelea* (e.g. Wolfe 1973). Reliable fossils of other genera within Juglandaceae have not been confirmed prior to the Teriary but nonetheless appear close in time to the Cretaceous-Paleogene boundary (Manchester 1987). Thus, we consider it highly likely that the divergence between *Rhoiptelea* and other Juglandaceae genera occurred prior to the end of the Cretaceous, and set 70 Mya as a minimum age for this node. We specify a modified uniform prior on the node, with 2.5% of the prior mass below the minimum and similarly 2.5% of the mass above the maximum bound.

2. MRCA of (Juglandoideae, Engelhardiodeae)
   Min: 65 Mya (hard bound)

**Justification:**

Fossils of *Polyptera manningii* and *Cyclocarya brownii* are recorded from the Paleocene (Manchester and Dilcher 1982, 1997; Lyson *et al.* 2019). Both are known from the Fort Union Formation, which has associated radiometric date estimates ranging from 61.03-64.73 Mya (Belt *et al.* 2004). Both of these taxa are members of Juglandoideae, judged on the basis of well-preserved vegetative and floral structures (Manchester and Dilcher 1997; Manos *et al.* 2007). Similarly, the earliest known fossil fruit assigned to *Cyclocarya* is recorded within a stratigraphic series encompassing the first 1 million years after the K-Pg mass extinction event with multiple associated radiometric dates Lyson *et al.* (2019). We thus set 65 Mya as a hard minimum bound for the divergence between Juglandoideae and Engelhardiodeae, specifying a truncated Cauchy prior distribution in MCMCTree with a heavy right tail.

3. MRCA: ((*Juglans*, (*Pterocarya*, *Cyclocarya*)), *Platycarya*)
   Min: 47 Mya

   **Justification:**
   The fossil genus *Cruciptera* from Eocene and Oligocene sediments of Western North America has been assigned with reasonable confidence to the crown group of Juglandeae (extant *Juglans*, *Pterocarya*, *Cyclocarya*) (Manchester 1991; Manos *et al.* 2007) on the basis of morphology. Radiometric dating of associated formations range from 43 to 47 million years old (Manchester 1991). We set this as the minimum age constraint for the stem age of Juglandeae (divergence from *Platycarya* using a truncated Cauchy distribution with a hard lower bound at 43 Mya and a heavy right tail.

4. MRCA of (*Cyclocarya*, *Pterocarya*)
   Min: 65 Mya (soft bound)

   **Justification:**

   *Cyclocarya* possess distinctive radially symmetric winged fruits, which are recognizable within the first one million years after the Cretaceous–Paleogene mass extinction (Lyson *et al.* 2019). However, as reliable fossils of *Pterocarya* are not known prior to the Oligocene (30 million years later than the earliest records of *Cyclocarya* fruits), we considered it plausible that early *Cyclocarya* fossils could represent stem lineages of extant *Cyclocarya* and *Pterocarya*. To reflect this uncertainty, we used a truncated Cauchy distribution with a 'soft' minimum bound at 65 Mya, a heavy right tail, and a mode at approximately 66 Mya. We allowed 20% of the prior mass for this node to fall below 65 Mya using a decaying exponential function over younger possible ages. Thus, the prior still strongly favors an older divergence for these taxa, as implied by the current taxonomic treatment of fossils assigned to *Cyclocarya*.

5. *Juglans* crown age
   Min: 28 Mya
   Max: 56 Mya

   **Justification:**
   Manchester (1987) reviews multiple fossil fruits attributable to sections *Rhysocaryon* (black walnuts, e.g. *J. siouxensis*) and section *Cardiocaryon* (butternuts, e.g. *J. lacunosa*) from the Early to Mid Oligocene. The divergence of these two sections represents the deepest divergence within the crown group of *Juglans*. *Juglans hindsii* represents a member of the black walnuts, while *J. mandshurica* represents a member of the butternuts. We therefore consider that the divergence of these sections occurred prior to the middle of the Oligocene period, and set a minimum age constraint for the MRCA of extant *Juglans* at 28 Mya. The first unambiguous *Juglans* fossil fruits are from the Mid Eocene, so we set the start of the Eocene as a maximum age constraint for this node.

We used a gamma distribution for the prior on this age, numerically finding the shape and scale parameters such that the prior mean was placed at 42 Mya, 2.5% of the prior mass was below the lower bound, and 10% of the prior mass was above the upper bound.

6. *Carya* crown age
   Min: 23 Mya Max: 56 Mya

   **Justification:**

   As discussed in (Manchester 1987), pollen of *Caryapollenites* appears in the Upper Paleocene and is similar to that of modern *Carya*, however its comparatively smaller size, together with the absence of definitive *Carya* macrofossils, suggest the crown *Carya* emerged after the Paleocene. We thus set the end of the Paleocene as an upper bound on this node. *Carya* is traditionally divided into four sections (*Carya*, *Apocarya*, *Sinocarya*, and *Ramphocarya*), with extant members of the former two distributed in North America, and extant members of the latter two distributed in East Asia. Molecular phylogenies have supported monophyly of North American and East Asian clades (Zhang *et al.* 2013, 2024c; Groh *et al.* 2025). Manchester (1987) reviews several fruit fossils which bear close affinity to recognized sections. For example, *C. quadrangula*, known from the Middle Oligocene appears to represent section *Apocarya*, while *C. strychnina*, known the Upper Oligocene, appears to represent section *Sinocarya*. The presence of fossils assigned to these two sections by the end of the Oligocene implies a divergence before the end of the Oligocene, so we set 23 Mya as a lower bound on this node. We specified a uniform prior between the minimum and maximum with soft bounds (2.5% prior mass below the lower and above the upper bound).

7. MRCA: *Pterocarya* sect. *Pterocarya*, *Pterocarya* sect. *Platyptera*
   Min: 5.3 Mya
   Max: 56 Mya

   **Justification:**
   Traditional classification of *Pterocarya* (Manning 1978) groups species into two sections, *Pterocarya* (e.g. *P. stenoptera*) and *Platyptera* (e.g. *P. macroptera*). These two sections were recently supported as monophyletic in what we consider the most comprehensive molecular phylogeny of the genus to date based on multiple nuclear loci (Song *et al.* 2020). Pollen similar to that of modern *Pterocarya* occurs in Eocene, and becomes more abundant through the Oligocene and Miocene Manchester (1987). There are isolated some reports of *Pterocarya*-like pollen from the Paleocene, but these warrant reinvestigation (Manchester 1987). Unambiguous fossil fruits of *Pterocarya* are not known prior to the Oligocene. Fossils that bear close affinity to the two modern sections (e.g. *P. smileyi*, sect. *Pterocara* and *P. asymmetrosa*, sect. *Platyptera*) are known from the Miocene (Manchester 1987). We thus specify a uniform prior for the age of this node, with soft lower and upper bounds between the start of the Eocene and the end of the Miocene.

### Supplementary Text S2

To investigate possible functional candidates within the G-locus, we used our denovo annotations of the *P. stenoptera* haplotypes, and also manually scanned alignments of RNAseq reads from all available tissues of *P. stenoptera* from this study and previous studies (Zhang *et al.* 2024d; Groh *et al.* 2025).

*GFAFL* is the only annotated gene that is conserved across all G-locus haplotype assemblies, but in addition there were single genes uniquely annotated in each of the *P. stenoptera* haplotypes. In the *G* haplotype, annotation predicted an additional 93 amino-acid open reading frame that was not present in the recessive haplotype annotation. A BLAST search revealed sequences with high similarity in variable numbers ($\sim$ 2-6 copies across *Pterocarya* assemblies), but no other assemblies showed similar sequence at the syntenic location on chromosome 11. These patterns could reflect transposition of an uncharacterized gene into the *P. stenoptera G* haplotype. We did not identify any known conserved domains within this predicted protein using the NCBI Conserved Domain search utility. RNA-seq reads from several tissues of both flowering types aligned here, and full-length IsoSeq transcripts from leaf tissue of *P. stenoptera* from (Groh *et al.* 2025), however these reads all contained several SNPs and a 2-bp indel that would render an early stop codon. These variants were not present in any of the homologous sequences identified in either haplotype assembly of *P. stenoptera*, suggesting these could potentially reflect mapping error from unassembled sequence or segregating presence/absence of paralogs within our mapping population. Nonetheless, the absence of this gene in the dominant haplotype of *P. macroptera* indicates that it is not conserved across heterodichogamous species, and the evidence of it encoding a functional protein is equivocal, so it is not likely to be the causal basis of the dimorphism in the ancestor of *Pterocarya*.

Turning to the expression data for the recessive haplotype of *P. stenoptera*, we investigated a predicted gene that was not annotated in the dominant haplotype. This predicted gene contains a single predicted exon with a 116 codon reading frame. A BLAST search identified homologs in *Juglans* and *Carya* annotated as 'glutathione S-transferase T3-like'. We did not see evidence of expression of this gene in any available tissues from *P. stenoptera*. Using BLAST we identified orthologous sequence in both the dominant and recessive G-locus haplotypes of *P. macroptera*, and the recessive haplotype of *P. fraxinifolia*. However, each of these contained stop codon mutations when translated, suggesting a lack of conserved function. This and the lack of evidence for expression in floral tissue suggests this sequence is not likely the causal basis of the flowering dimorphism.

Finally, using IsoSeq transcripts of *P. stenoptera* from Groh *et al.* (2025), we found compelling evidence for an additional gene within the recessive haplotype of *P. stenoptera*. IsoSeq transcripts were detected in two replicate samples of bark tissue, but not in floral tissue. A BLAST search identified homologs across eudicots, monocots, and magnoliids with the annotation 'AtMg00810-like' based on homology to an Arabidopsis gene. Closely homologous sequences were detected in over 10 copies in each *Pterocarya* assembly, but notably none were found within the respective G-locus regions. Thus, this gene is also an unlikely candidate for the control of heterodichogamy.

In contrast, *GFAFL* shows a conserved reading frame across all *Pterocarya* G-locus assemblies, and shows clear expression in male floral buds and mature male catkins of *P. stenoptera*. In RNAseq data of male catkin buds of *P. macroptera*, we detected a single sequencing read aligned to the *GFAFL* ortholog (out of 43.8 million read pairs). We note that this particular sample was collected in late November, well after the growth season, and expression of *GFAFL* appears to be reduced in dormancy in *Cyclcoarya* (Fig. 4c), and expression levels in male buds in *P. stenoptera* collected in late summer were somewhat low to begin with. Similarly, we did not detect expression of *GFAFL* in dormant male buds of *Cyclocarya paliurus* collected on the same day. However, we reanalyzed publicly available expression data from male and female floral buds of *C. paliurus* at the time of bud break (Chen *et al.* 2019) and detected clear expression of *GFAFL* in both tissues (we later determined this represented expression of two divergent alleles, see main text). We also note that *GFAFL* is clearly expressed in male floral tissue of two *Juglans* species (Fig. S10). Thus, *GFAFL* appears to be the strongest functional candidate within the G-locus for regulating heterodichogamy.

### Supplementary Figures

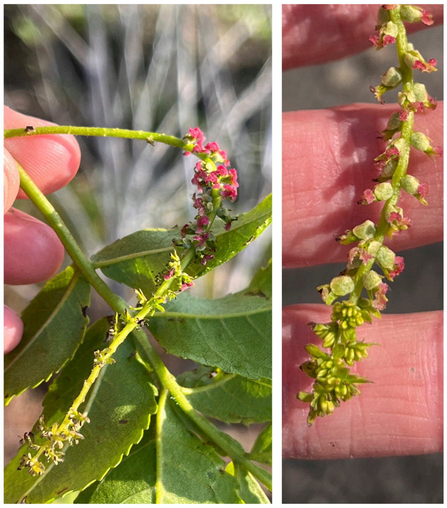

Figure S1: Examples of bisexual inflorescences of *Pterocarya stenoptera* uncommonly observed during the course of this study (inflorescences are typically unisexual). Male flowers are borne at the distal end of the rachis while female flowers (visible by pink stigmas) are borne near the proximal end. The inflorescences at left and right show protandrous and protogynous flowering, respectively. Photos by J.S.G.

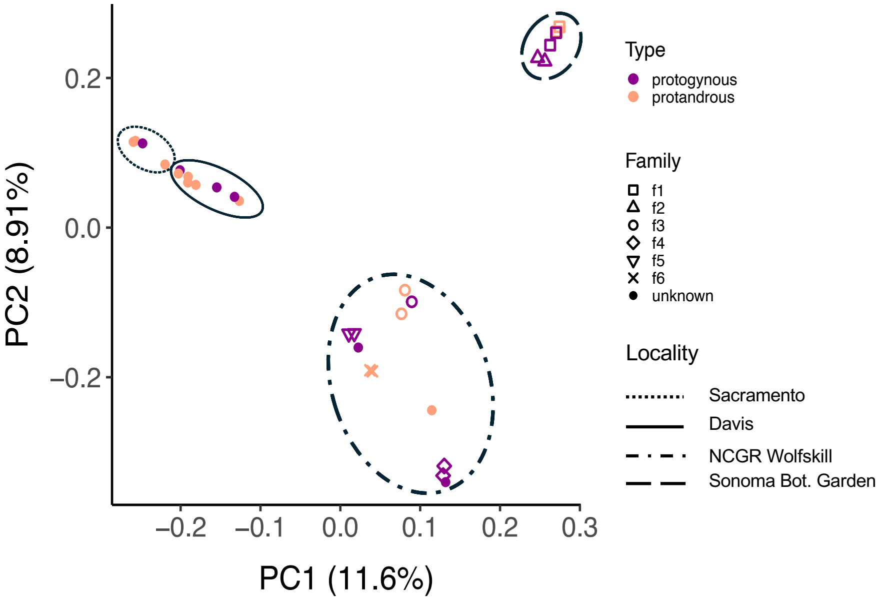


Figure S2: Principal Components Analysis (PCA) of genome-wide SNP variation in our mapping population of *P. stenoptera*. Families here are sets of offspring from the same maternal tree.

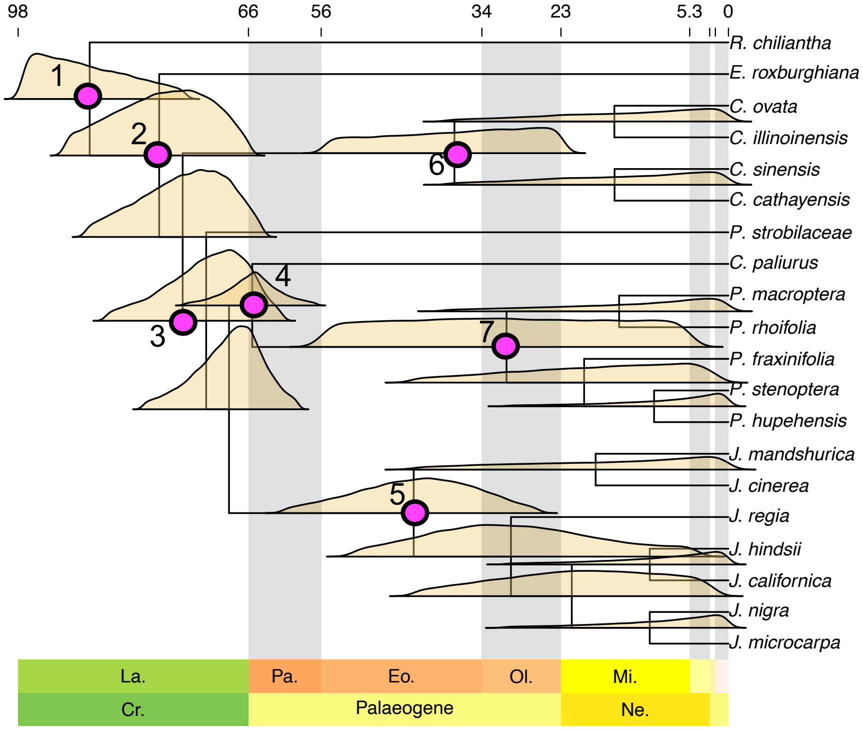


Figure S3: Visualization of the joint prior distribution on divergence times used in the MCMCTree analysis. Prior distributions were manually specified for a subset of nodes based on fossil information, and the joint prior on all nodes is subsequently generated by a birth-death model of speciation in MCMCTree. Nodes with manually specified priors are indicated in pink. Numbers correspond to the list of calibration points given in Supplementary Text 1. Note that the marginal prior for a given node may deviate from its specified distribution due to constraints of the birth death model that is used to specify the joint prior.

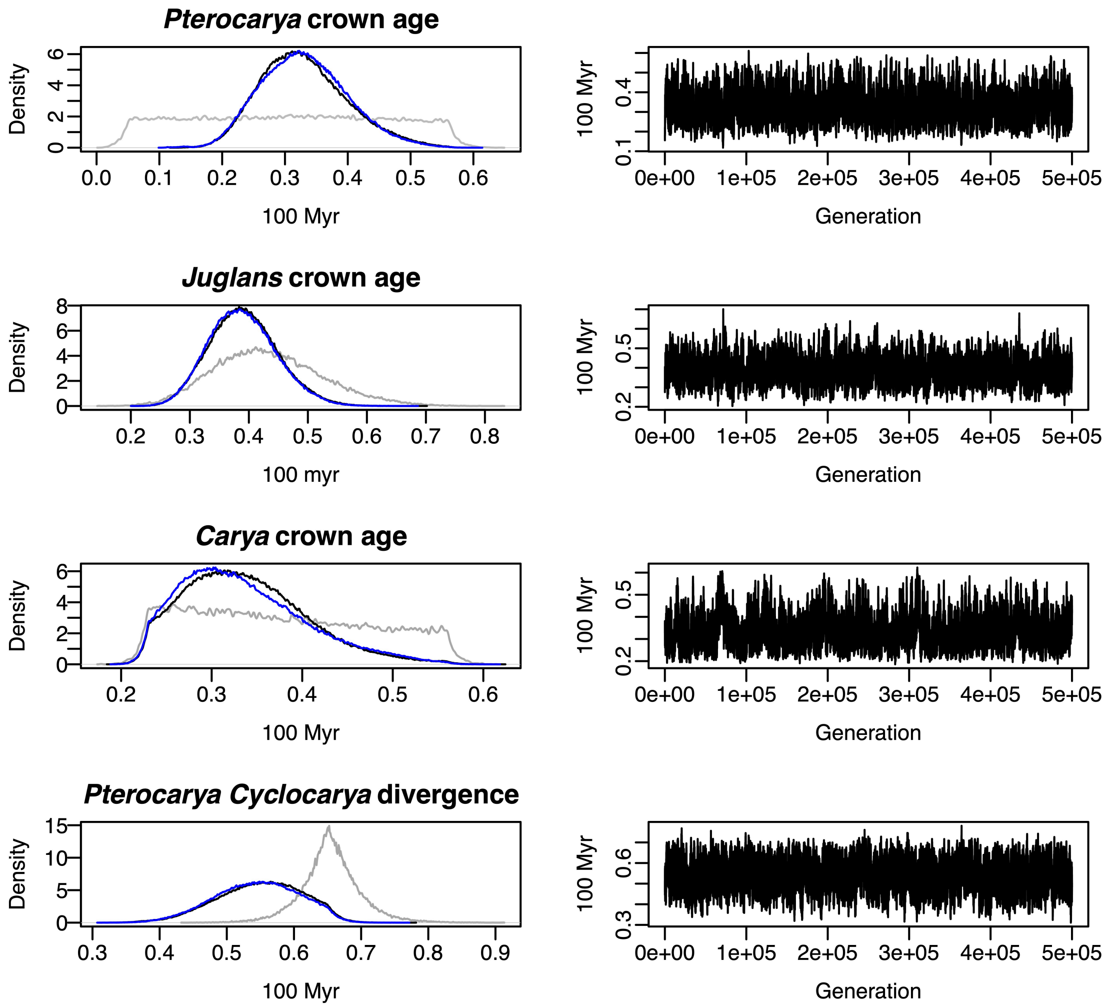


Figure S4: Diagnostics of Bayesian estimation of divergence times in Juglanceae using MCMCTree. We show the marginal prior density (from sampling, in gray) vs. marginal posterior density (black and blue show two chains run separately) and traces of a single MCMC chain for four nodes of interest. The priors for *Juglans* crown age and *Cyclocarya, Pterocarya* divergence were manually set, while the priors for *Pterocarya* crown and age and *Carya* crown age are derived from the birth-death model within MCMCTree. The molecular data appear highly informative for a more recent divergence between *Cyclocarya* and *Pterocarya* than what is reflected in the prior.

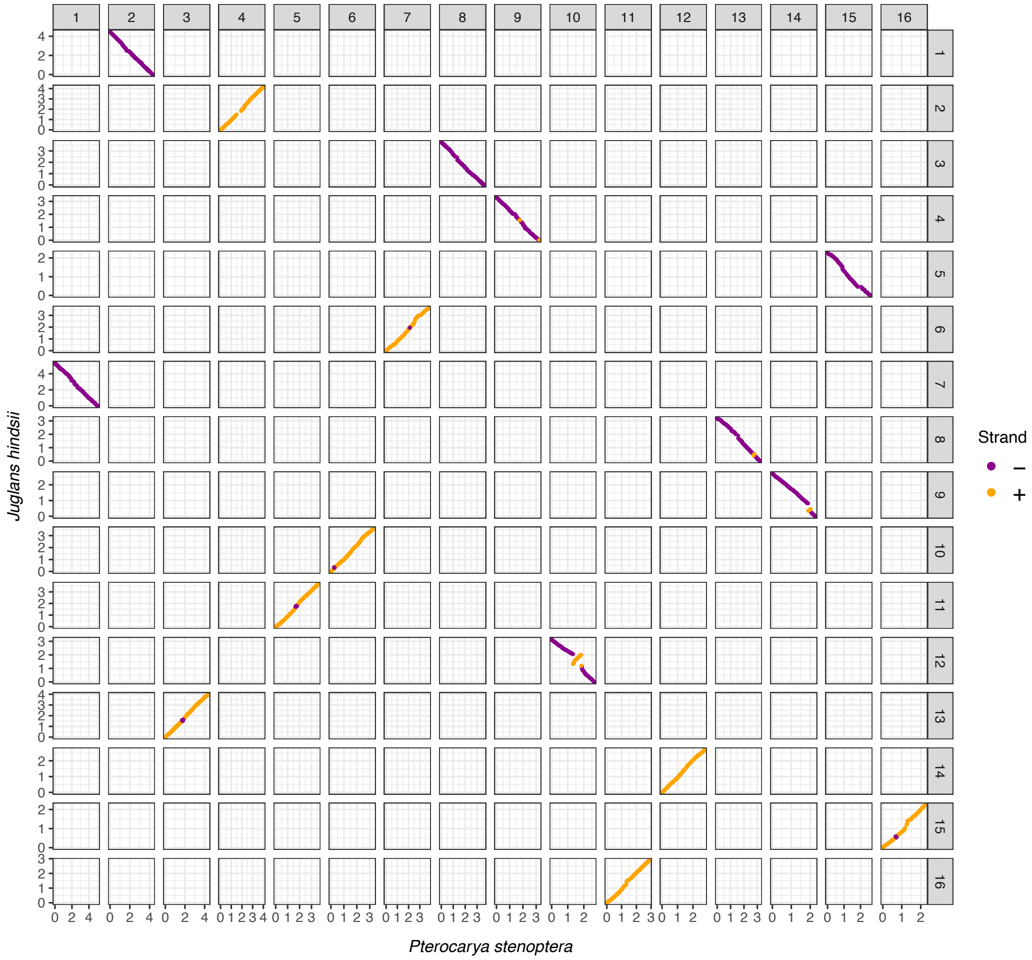


Figure S5: Chromosome-level synteny between our denovo assembly of *Pterocarya stenoptera* and an assembly of *Juglans hindsii* from (Groh *et al.* 2025). *P. stenoptera* was scaffolded by alignment to an assembly from the same species from (Zhang *et al.* 2024d) which incorporated Hi-C data, and chromosome names (shown in dark gray facet labels) match those of the reference used in scaffolding. The *J. hindsii* was reference-scaffolded to the high-quality assembly of *J. regia* which also incorporated Hi-C data, and follows the chromosome naming scheme for that species. The broad-scale 1:1 synteny between chromosomes reflects a known pattern of strong synteny conservation within Fagales. Axis units are 10 Megabases.

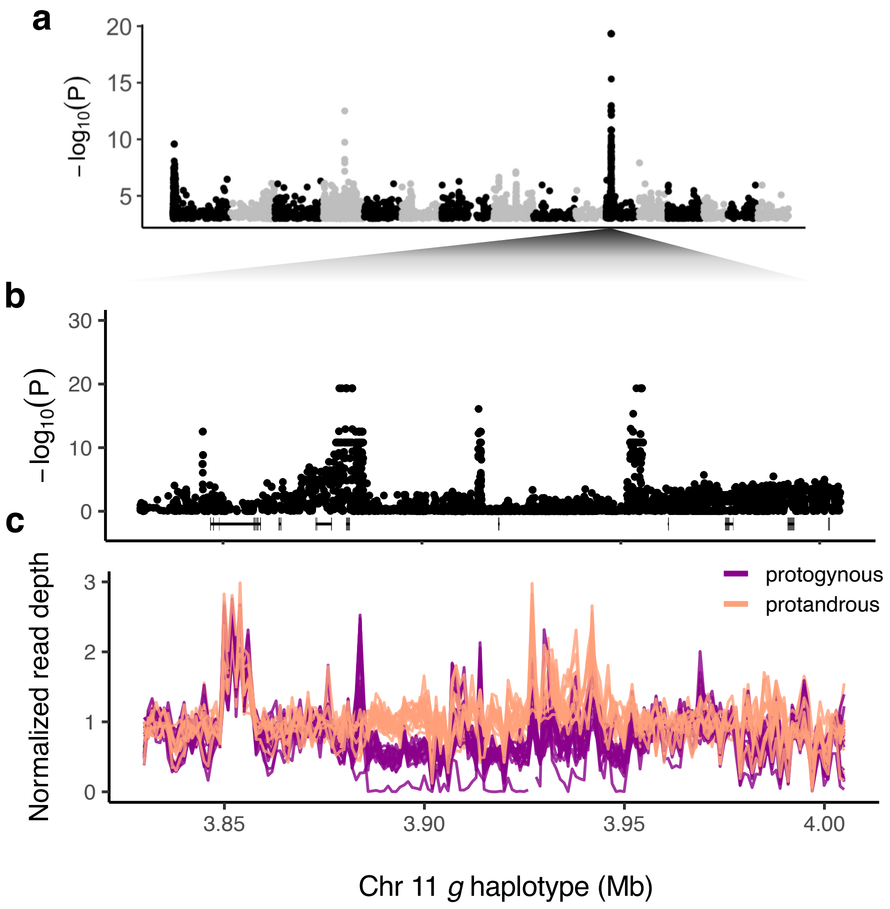


Figure S6: **a)** GWAS for dichogamy type using variants called against haplotype 1 of *P. stenoptera* (recessive) shows a strong peak on chromosome 11. The GWAS was considerably noisier in this case with many spurious peaks (vs. variant calls against haplotype 2, see Fig. 2). This discrepancy is typical for traits controlled by deeply diverged haplotypes with differing repeat content, and the results shown were obtained after masking repetitive regions. The y-axis is also truncated, hiding positions on chromosome 4 and chromosome 11 with perfect SNP associations. We confirmed that this extraneous peak on chromosome 4 represented alignment error; a BLAST search using the chromosome 4 local peak sequence as a query against the dominant haplotype matched a site within the GWAS peak in that haplotype. **b)** Zoomed in view of the GWAS peak with gene annotations at bottom. We found no evidence of expression for the predicted gene in the center of the associated region. **c)** Normalized average read depth in 1 kb windows for 30 phenotyped individuals. The individual with near zero read depth in the associated region was also identified as a *GG* homozygote on the basis of SNP genotypes and read depth against the *G* haplotype.

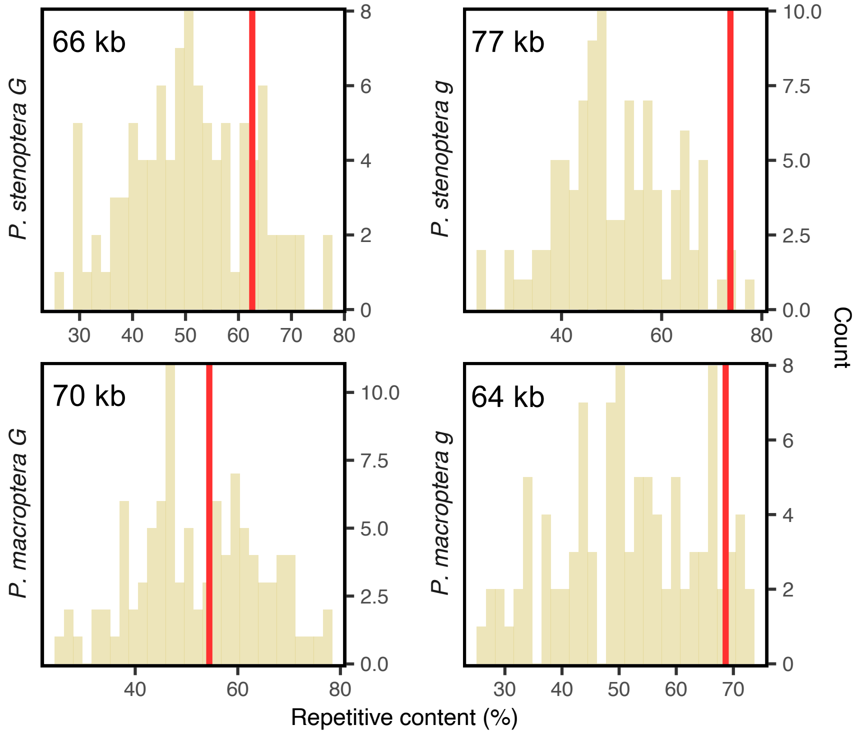


Figure S7: Repeat content of G-locus regions (red) compared with 100 neighboring genomic windows of equivalent size

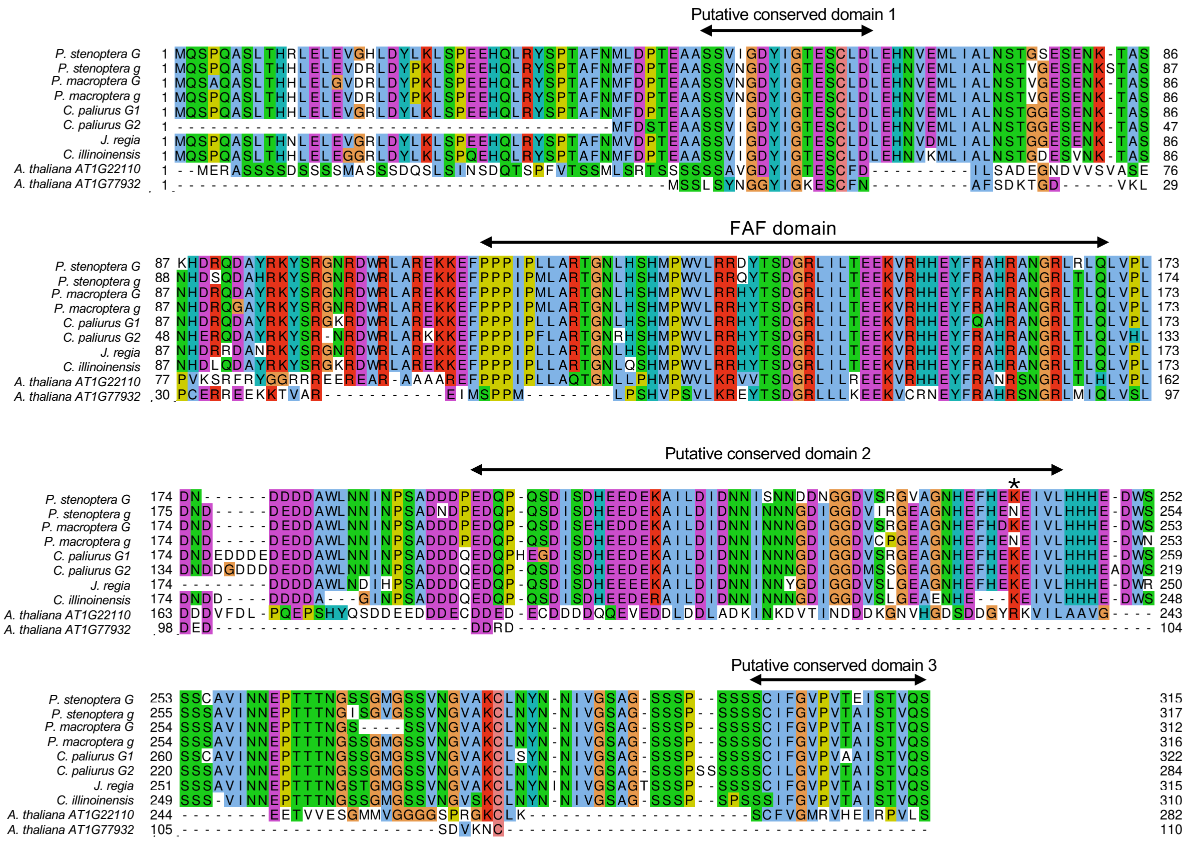


Figure S8: Amino acid alignment of *G-LOCUS-FAF-LIKE* orthologs from *Pterocarya Cyclocarya* and outgroups. We highlight four regions showing amino acid sequence conservation to the closest ortholog in Arabidopsis, one of which is the FAF domain. An asterisk marks the location of a nonconservative substitution within a conserved region which is the only nonsynonymous polymorphism between G-locus alleles shared across *Pterocarya*. At each site, amino acids are colored according to the conservation of amino acid properties in the sample acid properties using the default coloring scheme for Clustal X.

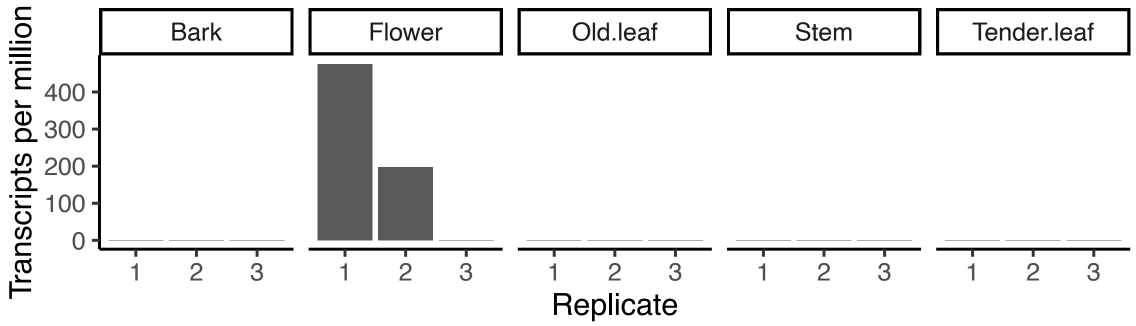


Figure S9: Analysis of the annotated ortholog of *GFAFL* in *Cyclocarya paliurus*. Raw data from (Zhang *et al.* 2024b). The original reference does not indicate whether the flower samples originated from pistillate or staminate flowers.

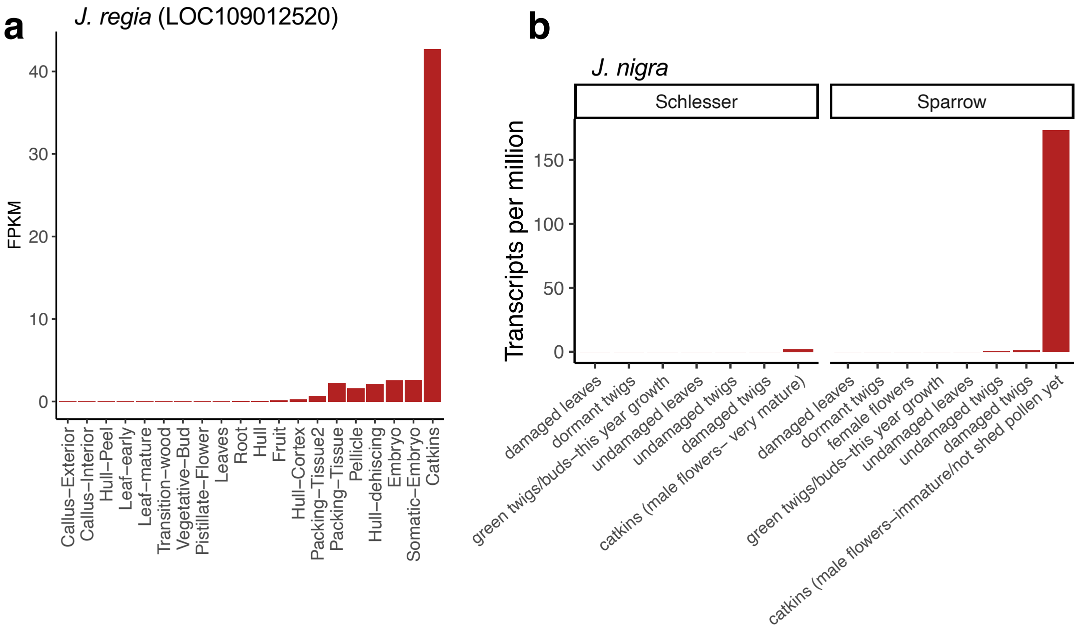


Figure S10: Expression of the ortholog of *GFAFL* in multiple tissues of two species of *Juglans*. **a)** Expression of *GFAFL* across 20 tissues sampled from *Juglans regia*. Here, catkins refers to the staminate inflorescences. Transcript quantifications were made available by Houston Saxe (UC Davis) (raw data from Chakraborty *et al.* 2016). Expression is measured as Fragments Per Kilobase of transcript per Million mapped reads. **b)** Expression of *GFAFL* across multiple tissues from two individuals of *J. nigra*. Raw data from (Hardwood Genomics Team 2015).

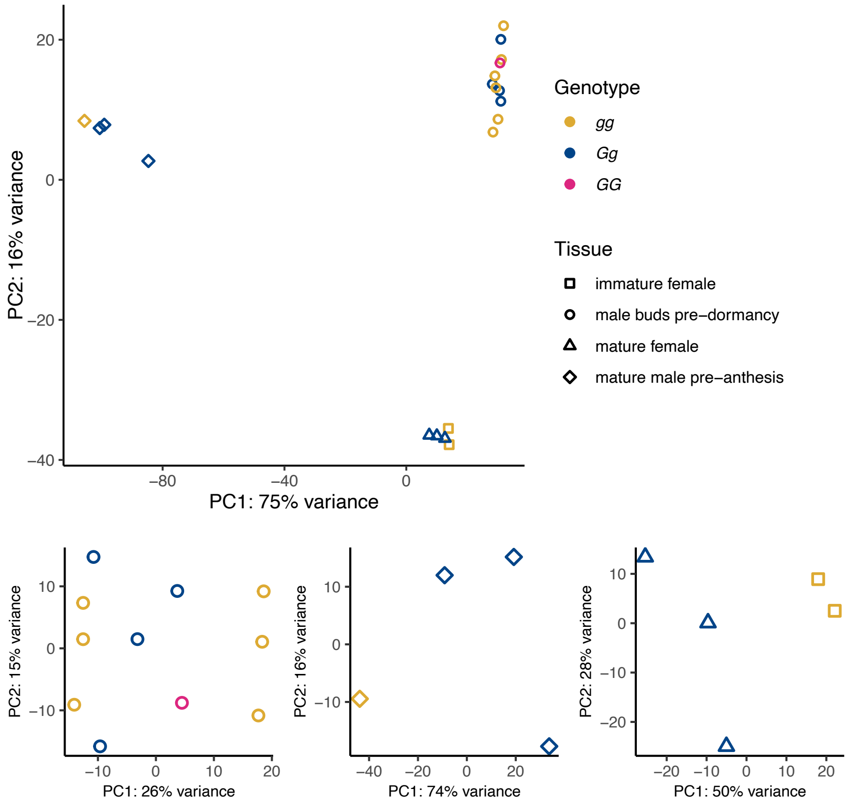


Figure S11: **Top)** Principal Components Analysis (PCA) biplots of variance-stabilized transcript counts for biological replicates across three floral tissue types of *P. stenoptera*. Within each tissue type, each point represents a different individual; some individuals were sampled for multiple tissue types. Samples cluster group by tissue type. Samples of female catkins and mature male catkins were sampled on the same day during the middle of the flowering season (March 20, 2024), while samples of pre-dormant male catkin buds were sampled on August 2 of the same year at the end of the growth season. **Bottom)** Separate PCA plots for each tissue run separately. We see evidence of clustering by morph the two floral tissues collected during flowering, albeit with low power to say whether this is significant.

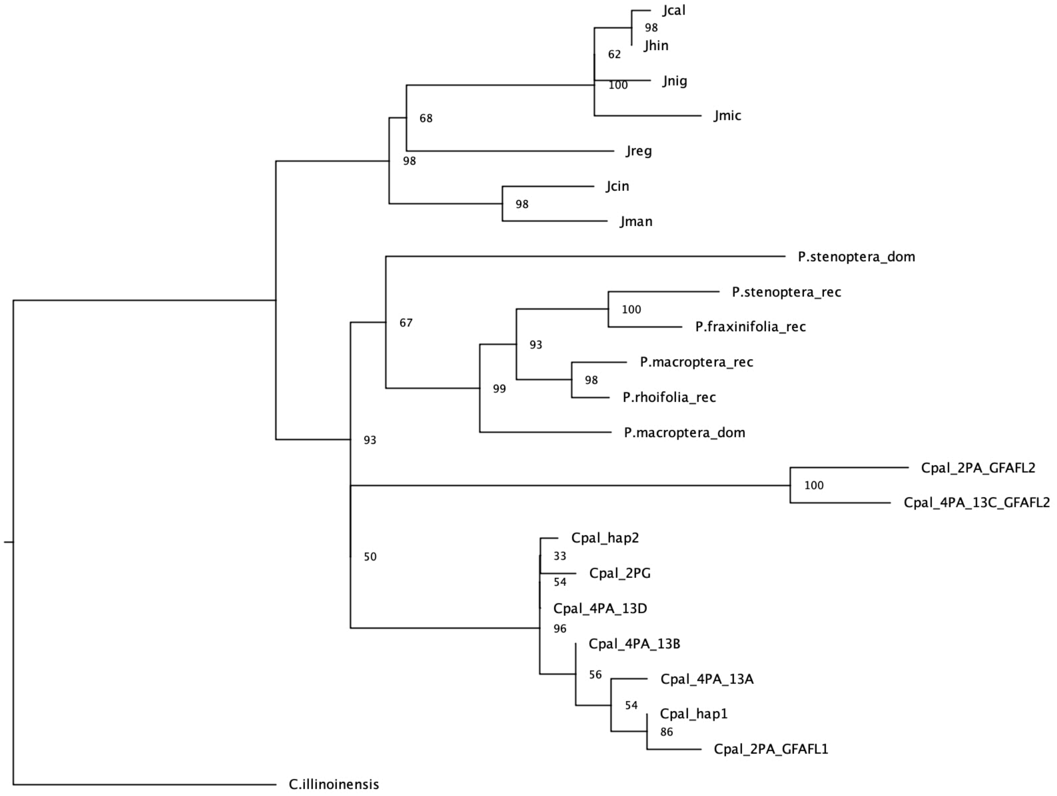


Figure S12: Maximum likelihood phylogeny of *GFAFL* orthologs in Juglandoideae constructed from an alignment of the coding sequence region.

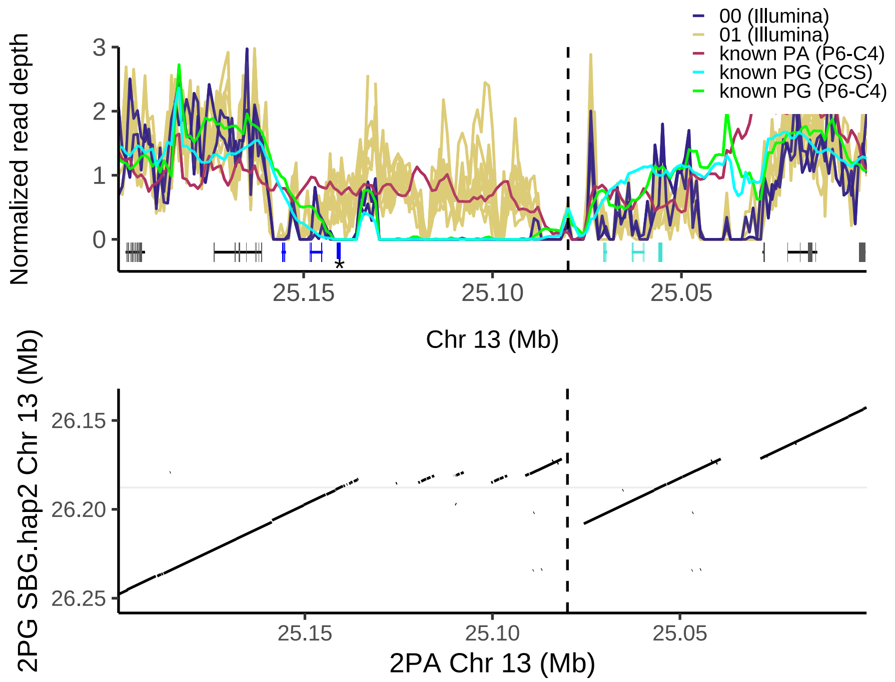


Figure S13: Read depth from 13 diploid *Cyclocarya paliurus* individuals against the diploid collapsed assembly from a protandrous *C. paliurus* individual from Qu *et al.* (2023a). This assembly appears to contain two tandem duplicated blocks of 3 genes each (turquoise and blue), including an annotated duplicate of *GFAFL* (blue, asterisk). However, we determined that the assembly in this region is chimeric, with two heterozygous haplotypes placed adjacent to another (see Supplementary Text 3), and these duplicate blocks are in fact orthologous single-copy alleles. The dashed vertical line indicates the inferred location of a contig break; no pacbio long reads from the individual used for assembly span this junction.

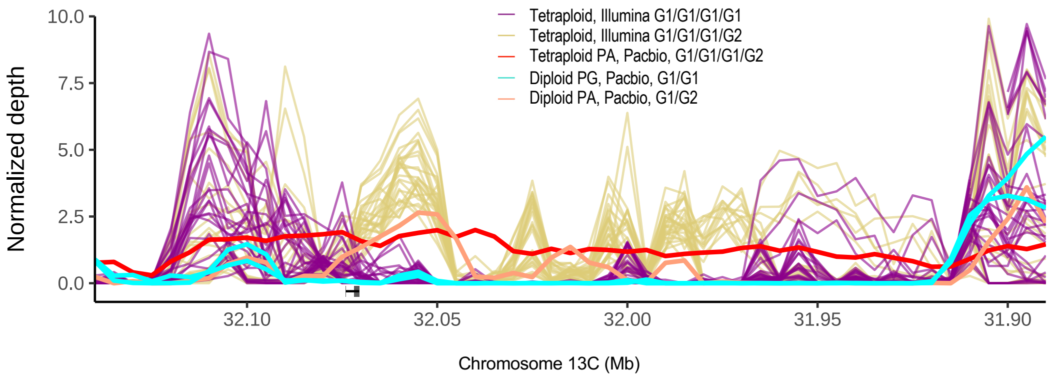


Figure S14: Read depth of *C. paliurus* individuals over the *G2* haplotype from a haplotype-resolved assembly from a protandrous tetraploid individual of the same species. The short read samples in tan and purple correspond exactly to the two tetraploid genotypic clusters shown in Fig. 4b, indicating that this copy number variant co-segregates with the *G2* copy of *GFAFL*. We also show sequencing depth from long-read Pacbio sequencing of four individuals of known type. The red line shows depth for the same individual used for assembly (Qu *et al.* 2023a). Blue lines correspond to our PacBio HiFi assembly of a diploid protogynous individual and the assembly reads for the diploid protogynous assembly from Qu *et al.* (2023a), the salmon line shows the protandrous diploid assembly reads from the same study. For each color, we indicate the inferred G-locus genotype.

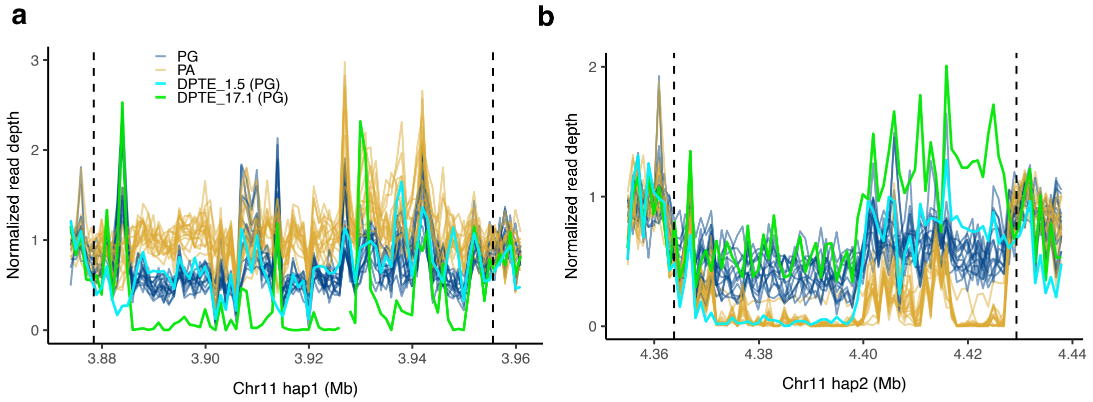


Figure S15: Read depth patterns of protogynous individuals suggest structural diversity of $G$ haplotypes. The individual DPTE_1.5 lacks depth across a portion of the $G$ haplotype in whole genome-resequence data, suggesting it may possess a $G$ haplotype that is structurally distinct from others in the sample. The sample DPTE_17.1, which we determine to be a $GG$ homozygote, shows a partial reduction in read depth across the same region, within the range of read depth seen in heterozygotes. This may indicate that one of its two $G$ haplotypes lacks this portion, however it could simply reflect poor read mapping to this region. We additionally note that one protandrous sample shows non-zero read depth across this region of the $G$ haplotype. We show elsewhere that this sample shows evidence of contamination on the basis of skewed allele depth ratios across this ratio (see methods and Fig. S20).

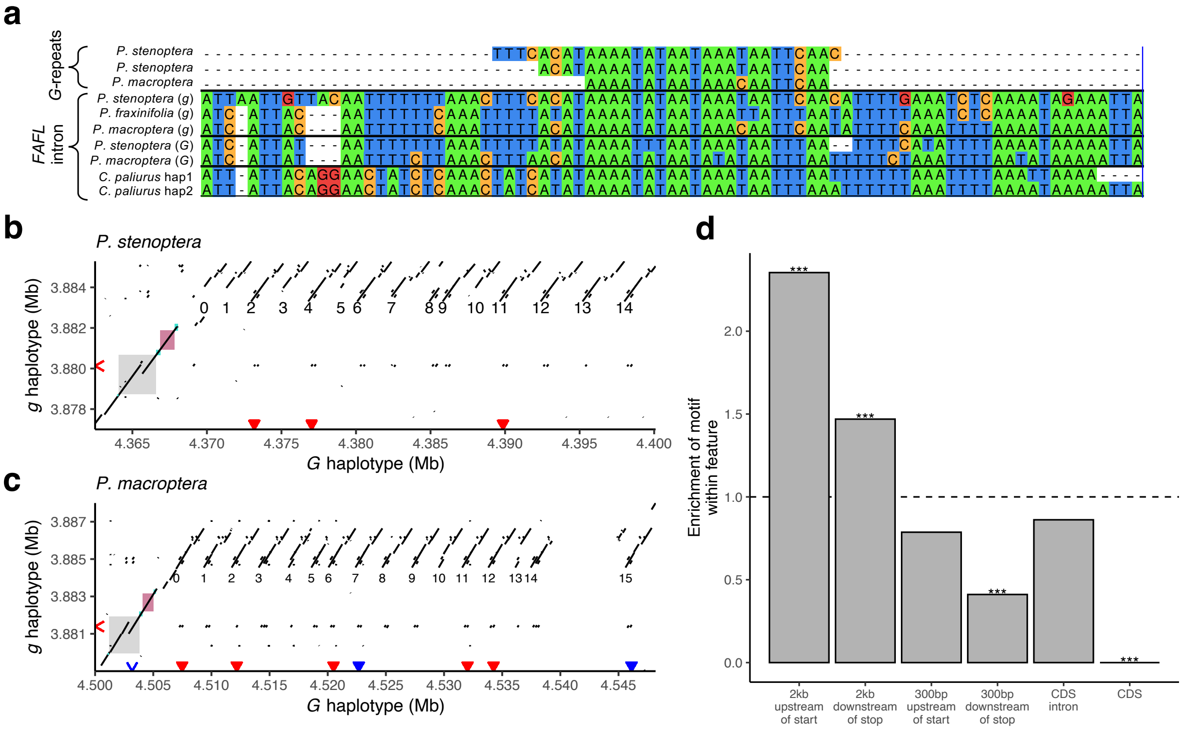


Figure S16: A candidate regulatory motif within $G$ haplotype repeats that may be associated with regulation of *GFAFL* expression. **a)** Alignment of an identified motif found in multiple locations within the $G$ haplotype insertion with the intron of *GFAFL*. Within both *P. stenoptera* and *P. macroptera*, the motif matches the $g$ but not $G$ intron. Consistent with a possible co-evolutionary relationship between these repeat motifs and the *GFAFL* intron, we identified SNPs in the sample where one allele is specific to the $G$ haplotype repeat and $g$ intron of *GFAFL*, separately in each species. **b,c)** Dotplot of alignments between $G$ and $g$ haplotypes in two species (shown for minimum alignment lengths of 21 bp and E value of 0.001). Colored arrows indicate positions where we identified matching of at least 21 bp between either intron and sequence anywhere within the entire G-repeat region. Shaded rectangles represent *GFAFL* coding sequence (maroon), intron (gray), and UTR (turquoise). **d)** The motif shown in (a) is enriched in regions proximal to genes, underrepresented in 3' UTRs, and entirely lacking in coding sequence, relative to the proportion of the genome contained within these features.

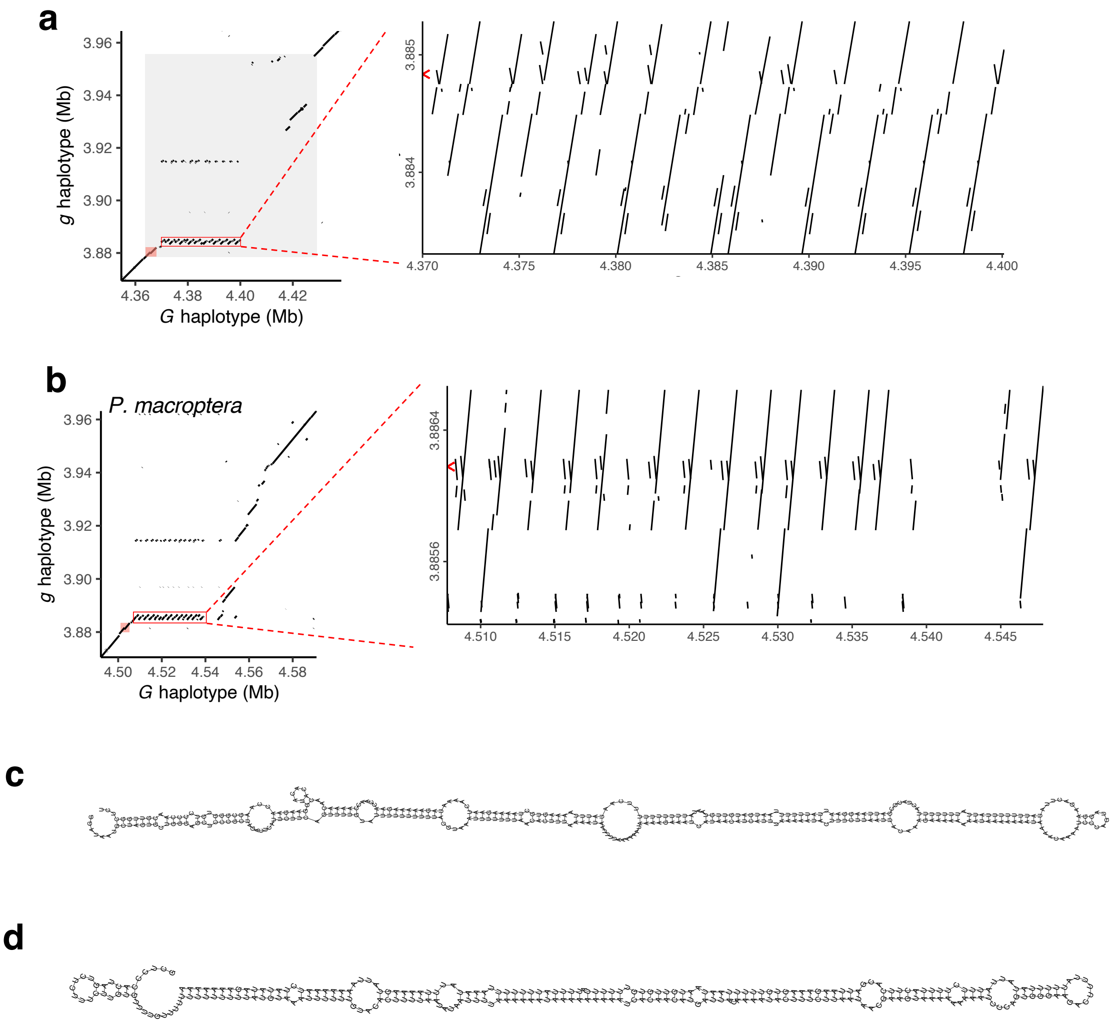


Figure S17: **a,b)** Pairwise alignments between the dominant and recessive G-locus haplotypes of two *Pterocarya* species reveal a shared inverted repeat region. At right, we show an inset of the regions highlighted in red rectangles. The red arrow in these plots indicates the coordinates in the recessive haplotype of a region with shared homology to the inverted repeat in both species. These regions in the recessive haplotypes are reciprocal best blast hits of one another, indicating that they are orthologous. **c,d)** Predicted hairpin RNA secondary structures from a selected inverted repeat in each species. (c) shows the predicted structure for repeat *G-7* of *P. stenoptera* and (d) shows the predicted structure for *G-0* of *P. macroptera*. Both of these predicted structures have a folding free energy of <0.2 kcal/mol/nucleotide, satisfying criteria for stability (Axtell and Meyers 2018).

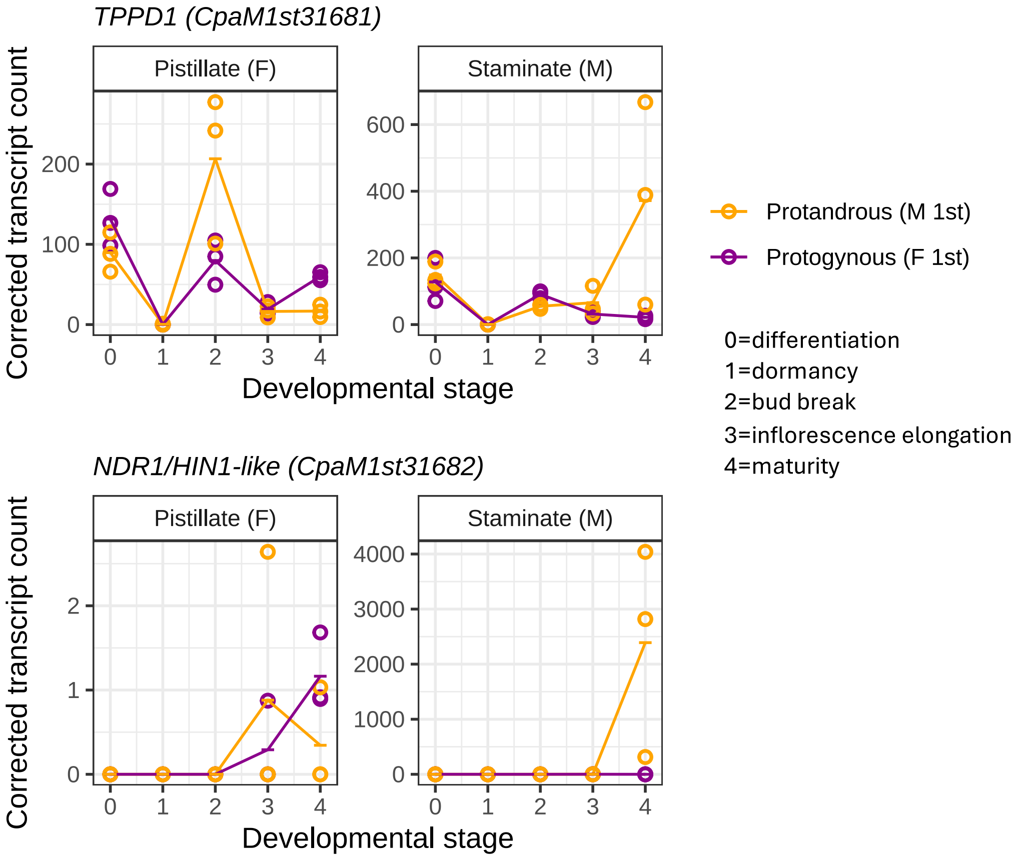


Figure S18: Expression of *Juglans* G-locus genes across a developmental time series dataset of gene expression from protandrous and protogynous morphs of *Cyclocarya paliurus* (Qu *et al.* 2023b). We note that the x-axis shows developmental stage rather than time, so some difference between morphs could reflect either sampling noise or expression changes that depend on time in the season rather than developmental stage.

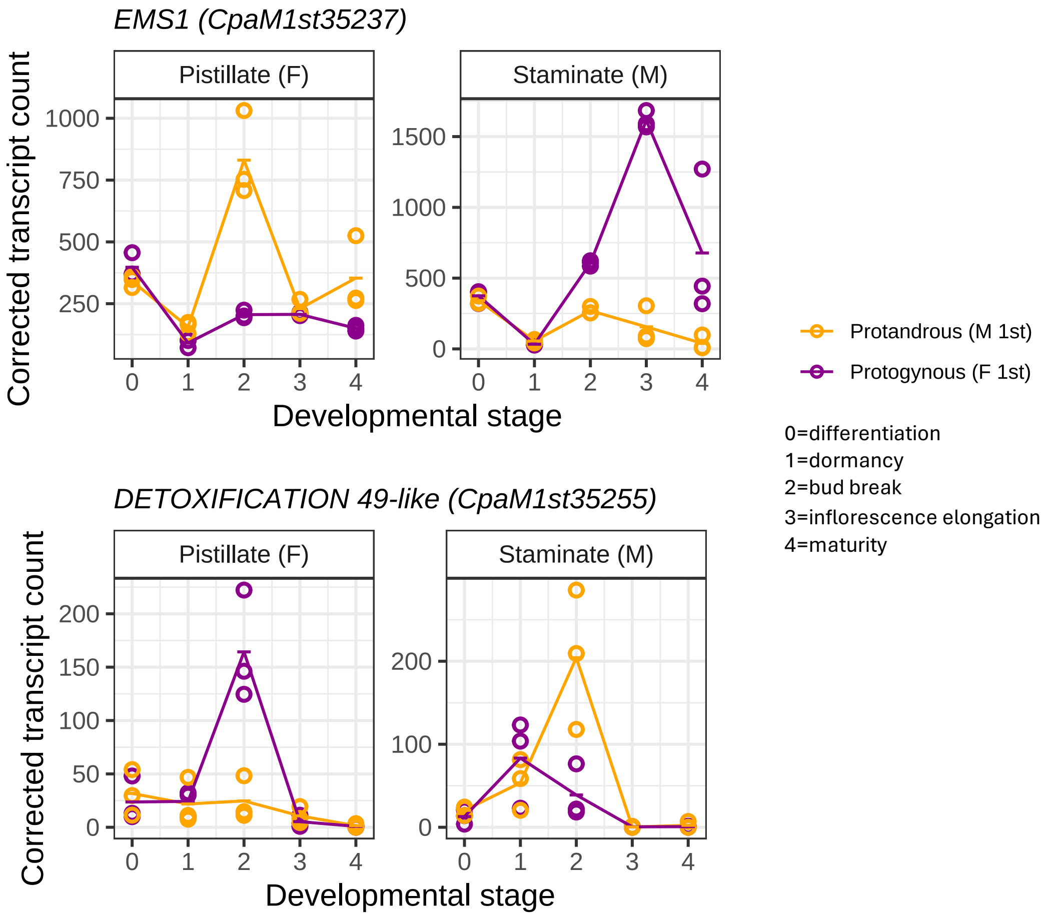


Figure S19: Expression of two candidate genes from the *Carya* G-locus across a developmental time series dataset of gene expression from protandrous and protogynous morphs of *Cyclocarya paliurus* (Qu *et al.* 2023b). These two genes show complementary patterns of expression between male flowers of in one morph female flowers of the other morph. These data suggest that, in *Cyclocarya*, *EMS1* could be negatively regulating flowering, while *DETOXIFICATION 49-like* could be positively regulating flowering.

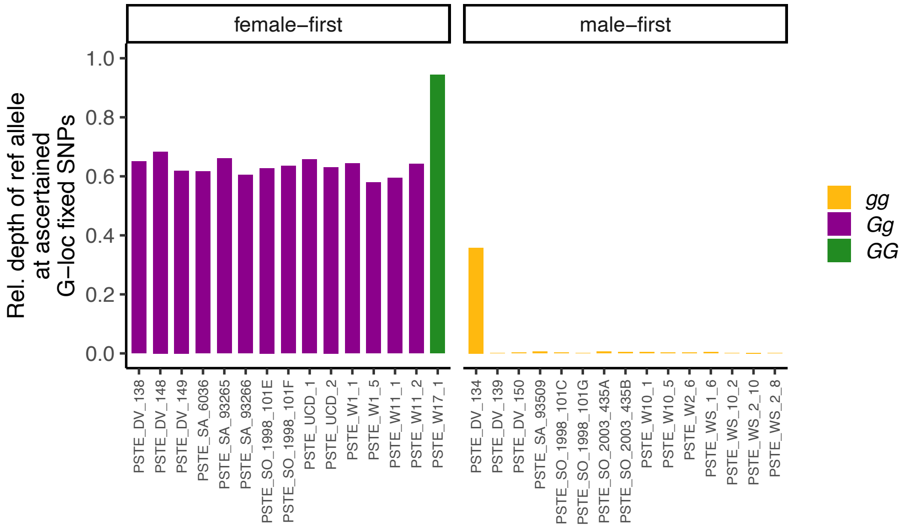


Figure S20: Allelic depths at a set of 255 SNP sites with fixed differences between G-locus haplotypes in *P. stenoptera*. One male-first sample (DV_134) was excluded in the ascertainment. We used this to investigate the possibility of this sample being chimeric as the basis for exclusion in analyses involving haplotype phasing. This sample showed non-zero depth of the reference ($G$) allele at many of these sites. Allele-specific read depth suggests this individual is not a heterozygote but rather was affected by sample contamination or the original sample contained material from two cell lineages with different G-locus genotypes. Given the deep divergence between haplotypes, heterozygotes should show a bias toward expression of the reference allele (here, $G$), which we do see for heterozygous protogynous samples (purple). In contrast, DV_134 shows the reverse bias in allelic depth, which is not consistent with this individual being a heterozygote. This individual was confidently phenotyped as male-first so we consider this pattern to reflect either a contaminated or chimeric sample and excluded it from a subset of analyses.
